# PERK inhibition blocks metastasis initiation by limiting UPR-dependent survival of dormant disseminated cancer cells

**DOI:** 10.1101/2021.07.30.454473

**Authors:** Veronica Calvo, Wei Zheng, Kirk A. Staschke, Julie Cheung, Ana Rita Nobre, Eduardo F. Farias, Ari Nowacek, Mark Mulvihill, Alan C. Rigby, Julio A. Aguirre-Ghiso

## Abstract

The unfolded protein response (UPR) kinase PERK has been shown to serve as a survival factor for HER2-driven breast and prostate cancers as well as for dormant cancer cells. However, its role in metastasis is not understood. Here we found in the MMTV-HER2 mouse model that quiescent HER2+ disseminated cancer cells (DCCs) displayed unresolved ER stress as revealed by high expression of the PERK-inducible GADD34 gene. *S*ingle cell gene expression profiling and imaging confirmed that a significant proportion of DCCs in lungs were dormant and displayed an active UPR. In human breast cancer metastasis biopsies, GADD34 expression and quiescence were also positively correlated. Importantly, PERK inhibition with a specific inhibitor (HC4) blunted metastasis development by selectively killing UPR^high^ quiescent but not proliferative DCCs. We also show that PERK inhibition altered optimal HER2 activity in primary tumors as a result of sub-optimal HER2 trafficking and phosphorylation in response to EGF. Our data identify PERK as a unique “Achilles heel” in dormant DTCs, supporting a requisite role for PERK in DTCs. Taken together, these data identify novel strategies to eliminate quiescent DCCs in patients with disseminated cancer disease.

## INTRODUCTION

Under stress conditions, the accumulation of unfolded proteins in the endoplasmic reticulum (ER) lumen activates three main pathways: PERK, IRE1α and ATF6. These pathways are part of a survival and adaptive mechanism known as the unfolded protein response (UPR) (Walter and Ron, 2011). Recent evidence suggests that in various types of cancer the UPR allows tumor cells to respond to increased demands on the ER and greater oxidative conditions imposed by an enhanced translational load caused by oncogenes, hypoxia, and other stress conditions (Blais et al., 2004; Chevet et al., 2015; Tameire et al., 2015). Oncogene-activated pathways increase ER client protein load by activating mTOR signaling and translation initiation (Hart et al., 2012; Ozcan et al., 2008; Tameire et al., 2015). Other studies have shown that PERK and the IRE1α-XBP-1 pathways contribute to the ability of cancer cells to adapt to hypoxia and microenvironmental stress (Bi et al., 2005; Blais et al., 2004; Chen et al., 2014; Romero-Ramirez et al., 2009; Rouschop et al., 2010; Schewe and Aguirre-Ghiso, 2008; Ye et al.), suggesting that the UPR enables critical adaptation mechanisms necessary to survive within a changing cellular milieu.

PERK activation initiates an antioxidant and autophagic response that coordinates a protection mechanism for mammary epithelial cells during loss of adhesion to the basement membrane (Avivar-Valderas et al., 2011). This survival response involves an ATF4 and CHOP transcriptional program (Avivar-Valderas et al., 2011) coupled to a rapid activation of the LKB1-AMPK-TSC2 pathway that inhibits mTOR (Avivar-Valderas et al., 2013). Human ductal carcinoma in situ (DCIS) lesions that displayed enhanced PERK phosphorylation, autophagy (Avivar-Valderas et al., 2011; Espina et al., 2010), and conditional ablation of PERK in the mammary epithelium had delayed mammary carcinogenesis promoted by the HER2 oncogene (Bobrovnikova-Marjon et al., 2010; Bobrovnikova-Marjon et al., 2008). Furthermore, HER2 increases the levels of proteotoxicity in tumor cells thereby activating JNK and IRE signaling and allowing HER2+ cancer cells to cope with this stress (Singh et al., 2015). Accordingly, 8 % of HER2-amplified human breast tumors display an upregulation of PERK mRNA, which further supports the notion that certain HER2+ tumors are dependent on PERK and/or other UPR pathways for survival (cBIOportal database (Cerami et al., 2012)).

We also reported that dormant (quiescent) cancer cells were dependent on PERK and ATF6 signaling for survival (Ranganathan et al., 2008; Ranganathan et al., 2006; Schewe and Aguirre-Ghiso, 2008). Furthermore, solitary quiescent pancreatic DCCs disseminated to the liver of mice also display a PERK-dependent UPR that was linked to loss of E-cadherin expression and downregulation of MHC-I, which favors immune evasion during dormancy (Pommier et al., 2018). In the MMTV-HER2 model, quiescent DCCs in bone marrow and lungs were also found to be E-cadherin negative (Harper et al., 2016), but the link to the UPR was not tested. Together, these data suggest that the UPR may serve as a stress and immune microenvironmental adaptive survival mechanism for DCCs.

Here we report that a previously described selective and potent inhibitor of PERK, HC4 (Calvo et al., 2021) can block HER2-driven breast cancer metastasis through the eradication of dormant DCCs. Imaging and single cell gene expression profiling revealed the existence of an UPR^high^/CDK inhibitor^high^ quiescent population of DCCs. In addition, CDK4/6 inhibition followed by HC4 treatment further decreased metastatic burden. Incidentally, PERK inhibition also prevented HER2-driven early cancer lesion development and induced stasis or regression of already established tumors *via* apoptosis. Our work reveals that PERK inhibitors, alone or in combination with anti-proliferative therapies, may represent a new strategy to target dormant cells during minimal residual disease stages and help prevent lethal metastases.

## RESULTS

### Quiescent HER2+ DTCs display an ER stress response

PERK pathway activation has been shown to serve as a crucial effector of UPR-induced growth arrest and survival linked to a dormant phenotype (Brewer et al., 2000; Ranganathan et al., 2006 and 2008). In the syngeneic HER2+ breast cancer model MMTV-HER2, a high percentage of mice develop metastases to the lungs, which can be initiated by early or late DCCs (Guy et al., 1992; Harper et al., 2016; Husemann et al., 2008). Dormant DCCs display loss of E-cadherin and expression of Twist1 (Harper et al., 2016) and E-cadherin-negative DCCs in pancreatic cancer models were also shown to be quiescent and displayed upregulation of CHOP, a PERK-induced gene (Pommier et al., 2018). We set out to determine whether in the MMTV-HER2 spontaneous metastasis model if this same correlation between levels of PERK pathway activation and cell cycle arrest existed. The correlation was evaluated by two different approaches, high resolution imaging using immunofluorescence (IF) and single cell resolution gene expression analysis of DCCs and metastasis. We performed IF of MMTV-HER2 lung tissue sections of animals bearing large tumors and thus bearing dormant and proliferative DCCs (Harper et al., 2016). Tissues were co-stained to detect DCCs positive for HER2, Ki67 (as a marker of proliferation) and GADD34 (or PPP1R15A). GADD34 is a PERK-inducible stress gene responsible for the programmed shift from translational repression (due to eIF2α phosphorylation) to stress-induced gene expression (Novoa et al., 2003). Image analysis showed that HER2+ metastatic lesions or solitary DCCs with a low proliferative index (ki67^low^) presented high levels of ER stress as shown by high levels of GADD34 expression (**Fig. 1a upper panels** and **graph**). On the other hand, highly proliferative DCCs or lesions showed very low levels of GADD34 staining (**Fig. 1a lower panels** and **graph**). The two markers, Ki67 and GADD34, were anti-correlated in 100% of the cells, supporting that UPR^high^ and quiescent DCCs and metastatic lesions can be identified via GADD34 detection.

**Figure 1.**
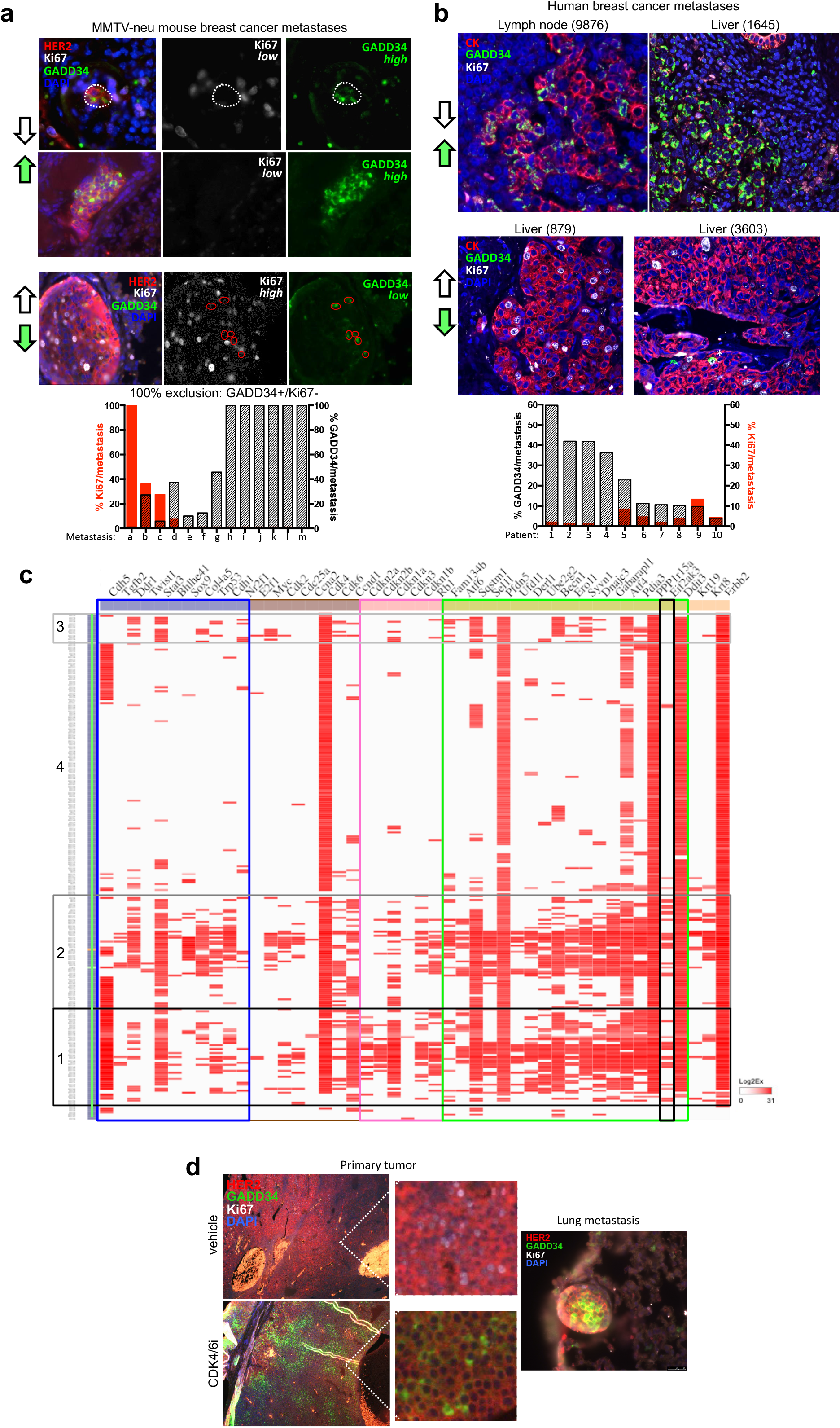
Quiescent disseminated HER2+ cells display high levels of ER stress PERK pathway activation. (a) Lung sections of MMTV-HER2 animals were stained for HER2, Ki67 (proliferation marker) and GADD34 (ER stress marker). The cells/met positive for either marker was quantified and shown as percentage of total cells (N=13). (b) Human breast cancer metastases from different locations (lymph node, liver, lung) were stained for cytokeratins, Ki67 (proliferation) and GADD34 (ER stress). The cells/met positive for either marker was quantified and shown as percentage of total cells (N=10). (c) Hierarchical clustering of the high-throughput targeted-gene expression (columns) profile of single cells (lung DTCs) (rows). Blue box, dormancy genes; brown box, cell cycle up genes; pink box, cell cycle down genes; green box, ER stress genes; black box, EIF2AK3 (PERK) gene. (d) Fluorescence IHC of tumor sections and lung sections from MMTV-HER2 females treated with Abemaciclib (50 mpk, 4 weeks) for HER2, Ki67 (proliferation) and GADD34 (ER stress). Scale bars, 100 µm.

We next tested if these correlations would also hold true in human breast metastatic lesions. HER2+ breast cancer metastases (n=10) to lymph node and additional 7 metastatic samples from different subtypes and tissues (lymph node, lung, liver) (**Supplementary Table 1**) were stained with a pan-cytokeratin cocktail to identify the metastatic lesions, Ki67 and GADD34. We observed that advanced human metastatic lesions displayed a more heterogeneous pattern of staining for both markers between different patients and in-between different areas of the same lesion than in the mouse model. However, a consistent negative correlation between levels of proliferation (Ki67) and ER stress activation (GADD34) was found, in HER2+ LN metastasis (**Fig. 1b**) or other target organs as well (**Supplementary Table 1**). This analysis validates the findings in the mouse models and that GADD34 may help identify UPR^high^/quiescent tumor cells in human metastatic sites.

Next, the analysis was expanded to markers of proliferation, quiescence, dormancy and ER stress in metastatic cells. To this end, we performed single cell targeted-gene expression analysis of DCCs, micro and macro-metastases lodged in lungs of MMTV-HER2 mice. Lungs from MMTV-HER2 females were processed into single cell suspensions and HER2+/CD45-cells were sorted (**Supplementary Fig. 1a**). The sorted cells were then processed for single cell separation, lysis, RT and pre-amplification using the C1 (Fluidigm) technology as shown in **Supplementary Fig. 1a**. This technique allowed us to isolate and process with a high degree of confidence (IF and molecular confirmation of HER2+ single cell) and quality 255 single DCCs and 90 primary tumor cells and their corresponding pools. Subsequently, high-throughput qPCR was used to analyze the expression of ER stress genes, cell cycle genes (both activators and inhibitors) and dormancy genes based on the literature (Kim et al., 2012; B’chir et al., 2013; Harper et al., 2016) (**Supplementary Fig. 1b**). The single cell resolution gene expression of DCCs revealed the existence of two populations of cells that are enriched for ER stress genes (groups 1 and 2, 41% of DCCs) (**Fig. 1c**), Groups 1 and 2. Group 1 (approximately 19% of the DCCs) showed concomitant and strong upregulation of all the ER stress genes tested (including PERK itself) (green box) with negative regulators of cell proliferation such as Rb1 and TP53 and CDK inhibitors p21, p27, p16 and p15 (pink box) (**Fig. 1c**). We also observed in these cells enrichment in the expression of dormancy genes such as NR2F1, DEC2 (*Bhlhe41*), TWIST1, CDH5, STAT3 and COL4A5 (Kim et al., 2012; Harper et al., 2016) (brown box). DCC group 2 (22%) also showed high levels of ER stress gene expression along with p21. In a third group (group 3 (6%)) ER stress, cell cycle inhibitors and dormancy genes were less prevalent, suggesting these might represent cells transiting out of dormancy or in cycling mode. In total, around 40% of the DCCs showed high to intermediate level of ER stress gene expression, concurrent with cell cycle inhibitors or dormancy genes. This is in range with the percentage of dormant DCCs detected in advanced progression MMTV-HER2 animals previously reported by our lab using phosho-Histone H3 and phospho-Rb detection (Harper et al., 2016). Taken together, these data illustrate that animals with detectable metastasis are comprised of ∼40% DCCs that display high expression of cell cycle inhibitor genes. Importantly, we further demonstrate in this model that dormant DCC subpopulations display an unresolved UPR with prominent expression of PERK pathway genes.

We have further correlated quiescence with heightened UPR through pharmacological inhibition of the UPR via our PERK inhibitor and other standard of care agents. UPR^high^ DCCs expressed higher levels of CDK inhibitors (**Fig. 1c**). Thus, we next asked whether treatment with a CDK4/6 inhibitor, Abemaciclib (50 mpk, 4 weeks), would result in decreased proliferation accompanied by an increase in UPR activation. Indeed, treatment of MMTV-HER2 females with Abemaciclib resulted in a striking increase in GADD34+ cells in primary tumor sections (**Fig. 1d**), which otherwise show very low and localized levels of GADD34 staining (*vehicle*). The increase in GADD34+ cells correlated with complete inhibition of proliferation as shown by Ki67 staining. An increase in GADD34 staining was also observed in lung metastases. This observation further supports the connection between induction of quiescence and UPR activation in primary tumor cells and disseminated cancer cells.

### PERK inhibition eradicates quiescent DCCs in bone marrow and lungs suppressing lung metastasis

The above findings opened the possibility of using selective PERK inhibitors to test whether inhibition of P-PERK could affect dormant DCC fate and metastasis formation. We used a PERK inhibitor derived from a 2-amino-3-amido-5-aryl-pyridine scaffold which we recently disclosed (Calvo et al., 2021). Briefly, HC4 was identified as a potent and selective PERK inhibitor with appropriate drug-like properties to support *in vivo* studies (Supplementary Table 7). HC4, along with other inhibitor variants from the amino-pyridine-mandelic acid-derived series (HC19, HC28), effectively decreased P-PERK (P-T980) levels in MCF10A cells expressing HER2 and in HEK293 cells stimulated with Tunicamycin (**Supplementary Fig. 1c**) as well as GADD43 induction upon thapsigargin treatment (**Supplementary Fig. 1d**) and rendered MCF10A/HER2 cells sensitive to low dose thapsigargin treatment. Taken together, these results showcase how selective inhibition of P-PERK with HC4 selectively affects adaptation to ER stress (**Supplementary Fig. 1e**). Using KINOMEscan™ kinase profiling (**Supplementary Tables 3-6**), HC4, HC19 and HC28 displayed high selectivity compared to other PERK inhibitors described in the literature and specifically GSK2656157 (Axten, 2017) even at a very high concentration (10 µM). The high specificity of HC4 for EIF2AK3 (PERK) over the eIF2α kinase family, EIF2AK1 (also known as HRI), EIF2AK2 (also known as PKR) and EIF2AK4 (also known as GCN2) or HER2 (**Supplementary Table 3**) indicates that the measured activity on eIF2α was highly specific to PERK inhibition.

Single dose administration of HC4 in CD1 female mice demonstrated that the compound is bioavailable following IP administration using ethanol/oil formulation and that the levels of HC4 in the plasma achieve a threshold that is well above that needed for PERK inhibition based on biochemical and cellular P-PERK IC50 values over a 24h time window (**Supplementary Fig. 1f**).

We treated 24-32 week old uniparous MMTV-HER2 female mice (which present an incidence of lung metastases of around 80%) with vehicle (see methods) or HC4 (50 mpk) IP daily, for two weeks, and collected mammary glands, lungs, pancreas, bone marrow and tumors for further analyses. HC4 was well tolerated, with no significant changes in body weight. The inhibitor did not have a significant effect on bone marrow cell homeostasis or on peripheral blood white cells as shown by no effect on total cell counts from MMTV-HER2 females (**Supplementary Fig. 1g**).

PERK inhibition caused a significant decrease in P-PERK and P-eIF2α levels in the mammary gland ducts and in pancreatic tissue (although only partial inhibition was observed at this dose, especially in pancreatic islets) (**Fig. 2a**). We conclude that systemic HC4 delivery effectively inhibits PERK activation and eIF2α phosphorylation. The inhibition of PERK did not fully deplete PERK activity, which may allow mice to control their pancreatic function and glucose levels (Yu et al., 2015). Moreover, in a separate study in mice dosed orally with HC4 for 28 days, where similar exposures were achieved to the 50 mpk IP dose noted above, no deleterious effects on the pancreas or with clinical chemistry (ie insulin and glucose levels) were observed (data not shown).

**Figure 2.**
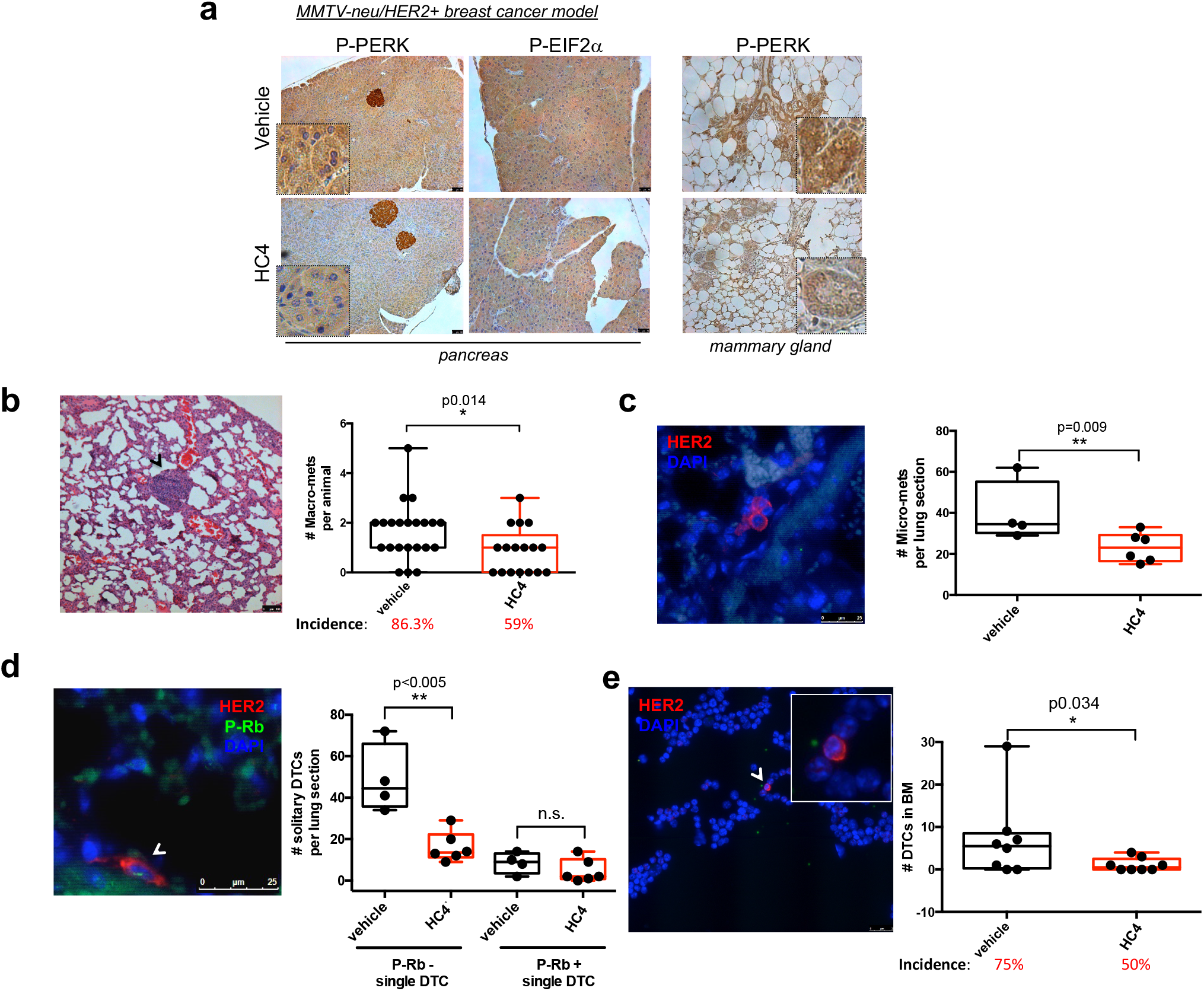
HC4 PERK inhibition decreases metastatic disease in lungs and bone marrow at the single disseminated cancer cell level. (a) MMTV-HER2+ females (24-week-old) were injected daily with vehicle or HC4 (50 mpk) for 2 weeks. Immunohistochemistry (IHC) of pancreas and mammary gland sections with antibodies to P-PERK and P-EIF2α. Inserts show higher magnifications. Scale bars, 100 µm. (b) Macro-metastases (>100 cells) were detected by H&E staining and quantified in 5 lung sections/animal (N=16). Scale bar, 100 µm. P by Mann-Whitney test. (c) Micro-metastases (2-100 cells) were detected by IHC staining using an anti-HER2 antibody and quantified per lung section/animal ± s.d. (N=6). Scale bar, 25 µm. P by Mann-Whitney test. (d) Solitary disseminated cancer cells (DCCs) were detected by IHC staining for HER2, classified as P-Rb+ or P-Rb-and quantified per lung section ± s.d. (N=6). Scale bar, 25 µm. P by Mann-Whitney test. (e) Disseminated cancer cells in bone marrow were detected by IF staining for CK8/18 and HER2 in cytospins from mature hematopoietic cell-depleted bone marrow tissue (N=8). Scale bar, 25 µm. P by Mann-Whitney test.

MMTV-HER2 animals develop metastases to the lungs, which can be initiated early in progression (Guy et al., 1992; Harper et al., 2016; Husemann et al., 2008). Thus, we next monitored the effect of HC4 on metastatic disease in animals with small and/or palpable large tumors. All the vehicle-treated animals presented metastases detectable in sections stained with H&E. Lesions that displayed >100 cells were categorized as macro-metastases as they are also commonly positive for proliferation markers (**Fig. 1a**). The quantification of macro-metastases per animal (5 non-consecutive lung sections) revealed that, after just a two-week treatment, HC4 reduced the number and the incidence of macro-metastases (**Fig. 2b**) while not affecting the area of these metastases (**Supplementary Fig. 2a**). This suggested that PERK inhibition through HC4 might be acting on the initial steps of metastasis rather than shrinking established macro-metastases. Thus, we tested whether HC4 treatment might be affecting the intravasation of tumor cells from the primary site or the transition from solitary DCC to micro-metastasis (containing 2-100 cells). Detection of HER2+ circulating tumor cells (CTCs) directly in blood samples showed no significant difference between vehicle and HC4-treated animals (**Supplementary Fig. 2b**), indicating that HC4 is not grossly affecting the intravasation of tumor cells. On the other hand, detection of micro-metastasis and single DCCs using HER2 detection via IHC revealed a significant decrease in the number of micro-metastases in HC4-treated females (**Fig. 2c**). More than 80% of single DCCs in lungs are negative for P-Rb, indicating that they are mostly out of cycle and dormant. This measurement reproduces observations noted in our previous publication (Harper et al., 2016). HC4 significantly reduced the number of non-proliferating (P-Rb negative) single DCCs that are commonly associated with blood vessels in lung sections, while not affecting the number of P-Rb positive solitary DTCs (**Fig. 2d**) or micrometastases (**Supplementary Fig. 2c**). Importantly, HC4 significantly decreased the number of DCCs found in bone marrow (**Fig. 2e**). In this organ, metastases never develop but DCCs are found at a high incidence and are dormant (Bragado et al., 2013; Husemann et al., 2008). These results argue that PERK inhibition is selectively targeting non-proliferative dormant DCCs that display active PERK and UPR signaling.

### PERK inhibition blocks HER2-driven early and late mammary primary tumor progression

Having demonstrated that there is a dependency on PERK in quiescent UPR^high^ DCCs, where dormancy is most relevant, we shifted our attention to primary tumor lesions. HER2-driven progression was found to be genetically dependent on the PERK kinase in the MMTV-HER2 model (Bobrovnikova-Marjon et al.) and a recent study showed that HER2+ tumors are sensitive to proteotoxicity and dependent on ERAD for survival (Singh et al., 2015). Further, cBIO database (Cerami et al., 2012) analysis showed that ∼14% of HER2-amplified human breast tumors (Breast Invasive Carcinoma, TCGA, Nature 2012 dataset) display upregulation of the mRNA for PERK (**Supplementary Fig. 3a**). Thus, we investigated whether HC4 affected HER2-induced breast tumor progression in primary lesions where the different stages of progression from hyperplastic mammary glands through DCIS and invasive cancer can be dissected (Lu et al., 2010; Muller et al., 1988).

Analysis of 24-week old uniparous female mammary glands showed that vehicle-treated MMTV-HER2 animals exhibited ducts with secondary and tertiary dense branching (**Fig. 3a,** left panels), and histological analysis showed frequent mammary hyperplastic lesions (**Fig. 3a,** right panels, black arrows). In contrast, HC4-treated animals showed a “normalized” glandular architecture with less dense branching, resembling the mammary tree of non-transgenic normal FVB mice (**Supplementary Fig. 3b**). HC4-treated animals also showed a dramatic increase in the number of hollow lumen mammary gland ducts, constituting more than 60% of the structures compared with around 20% in control females (**Fig. 3b** and **Supplementary Fig. 3c**). The number of occluded hyperplasias and DCIS-like lesions was also reduced to less than half of that observed in vehicle-treated animals. Hyperplastic lesions in control HER2+ animals showed varying degrees of luminal differentiation as assessed by the uneven levels of cytokeratin 8/18 expression (**Fig. 3c, upper panel**). The myoepithelial cells (detected as smooth muscle actin, SMA, positive), otherwise equally spaced in normal FVB animal ducts, were unevenly distributed in the vehicle-treated hyperplasias in the MMTV-HER2 mice. In contrast, HC4-treated MMTV-HER2 animals presented increased expression of cytokeratin 8/18 in the luminal layer, frequently surrounding an empty lumen, and an external continuous layer of myoepithelial cells (**Fig. 3c**, lower panel and graph). This data indicates that HC4 treatment leads to a restored differentiation state of early HER2-driven cancer lesions.

**Figure 3.**
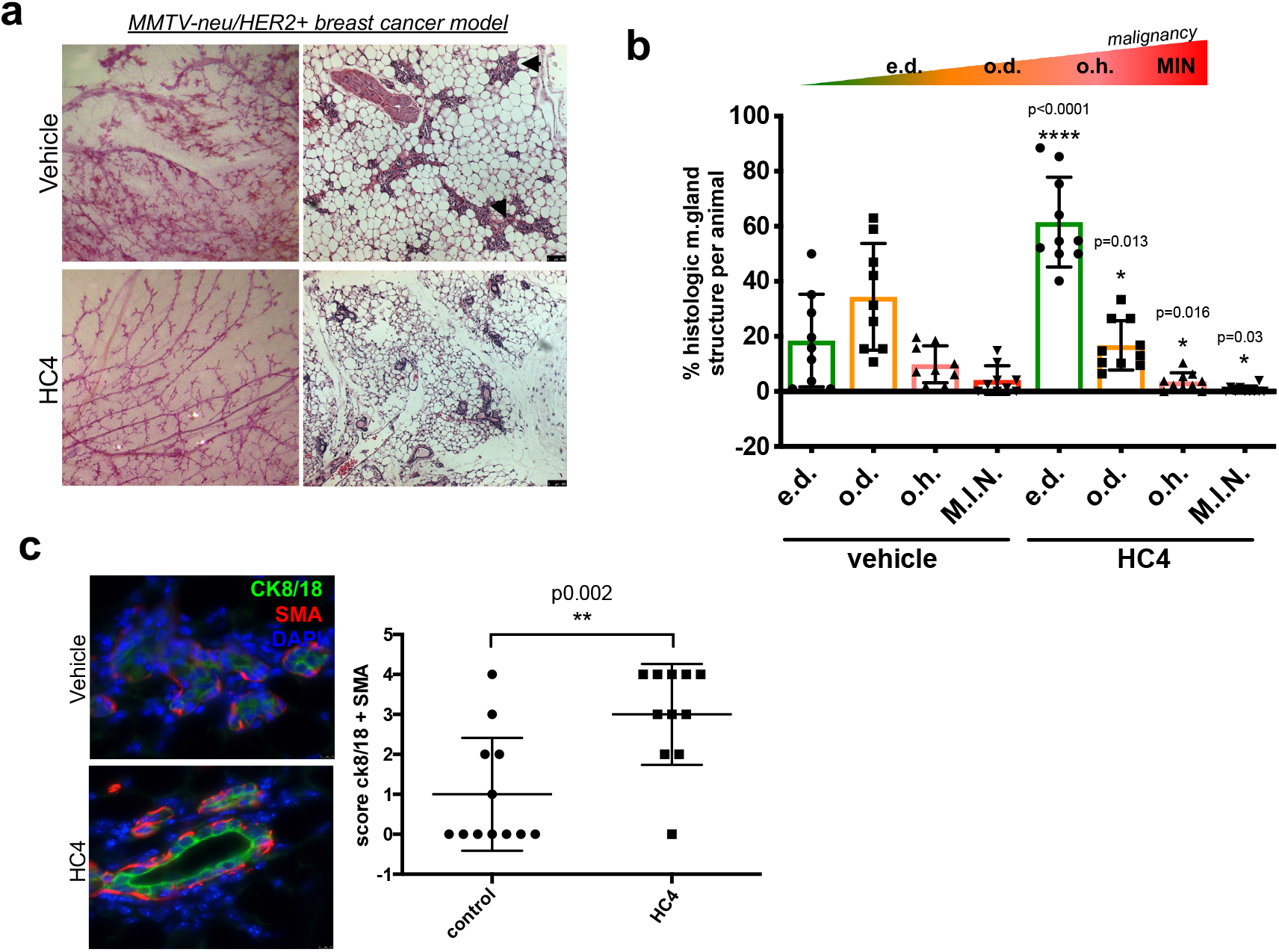
The PERK inhibitor HC4 causes mammary gland “normalization” in the MMTV-HER2+ breast cancer model. (a) Representative images of carmine staining of whole mount mammary glands and H&E-stained mammary gland sections from vehicle- and HC4-treated animals. Scale bar, 100 µm (b) Quantification of histological structures (empty duct e.d., occluded duct o.d., occluded hyperplasia o.h. and DCIS-like mammary intraepithelial neoplasia M.I.N) present in H&E-stained mammary gland sections (N=50/animal, animals N=13) found in vehicle- and HC4-treated animals ± s.e.m. Statistical significance (p) calculated by Mann-Whitney test. (c) IHC for epithelial luminal marker cytokeratin 8/18 (CK8/18) and myoepithelial marker Smooth Muscle actin (SMA) in mammary gland sections. Score for CK8/18+ and SMA+ structures per animal, N=12. P by Mann-Whitney test. Scale bar, 75 µm.

We next treated animals once they displayed tumors, ranging from 30 to 200 mm^3^ volume (two tumors were >200 mm^3^) for two weeks with HC4 (**Supplementary Fig. 4a**). In the vehicle treatment group, tumors grew steadily (**Fig. 4a**), reaching up to 10 times its original volume in two weeks (**Supplementary Fig. 4b,** upper graph). In contrast, HC4-treated tumors showed a reduced growth rate (**Fig. 4a**), with some tumors remaining in complete cytostasis (defined as doubling tumor volume only once in the 2-week period, 43% in HC4-treated vs 7% in controls) (**Supplementary Fig. 4b**, lower graph) and some tumors (25%) showing regression in the 2-week window treatment (**Supplementary Fig. 4c**). This led to a significant decrease in median final tumor volume (**Fig. 4b**). While the levels of proliferation (P-histone H3 IHC) were not different between vehicle-and HC4-treated tumors (**Supplementary Fig. 4d**), TUNEL staining of tumor sections showed a significant increase in the levels of DNA fragmentation present in HC4-treated animals (**Fig. 4c**). Thus, in overt primary lesions HC4 treatment induced apoptosis of established HER2+ tumors, arguing for context-dependent fitness-promoting functions of PERK during progression.

**Figure 4.**
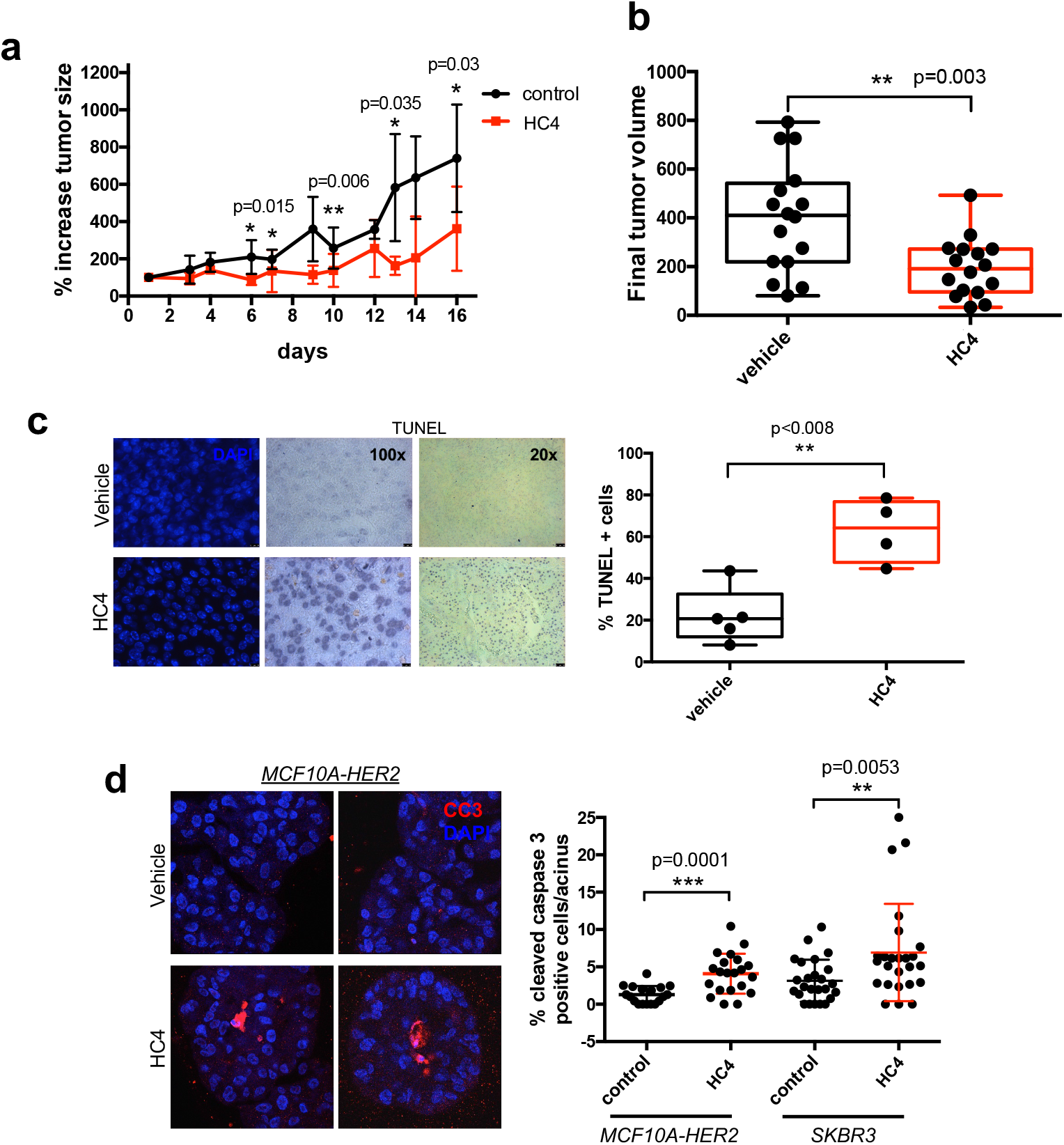
PERK inhibition impairs tumor growth in MMTV-HER2+ females. (a) MMTV-neu females (24- to 32-week-old) presenting overt tumors were injected daily with vehicle or HC4 (50 mpk) for 2 weeks. Percentage variation of tumor size in vehicle- and HC4-treated animals ± s.d. (N=16). P by Mann-Whitney test. (b) Final tumor volume (mm3). The whiskers represent the min and max of the data (N=16). P by Mann-Whitney test. (c) Representative IHC of TUNEL staining to measure apoptosis levels in tumor sections. Scale bars, 10 and 50 µm. Graph, percentage TUNEL positive cells in vehicle- and HC4-treated tumor sections (N=5). P by Mann-Whitney test. (d) HER2+ MCF10A-HER2 or SKBR3 cells were seeded on Matrigel and after acinus establishment (day 4) wells were treated with vehicle (control) or HC4 (2 µM) for 10 days. Percentage of cleaved caspase-3 positive cells per acini (N=20) ± s.d. P by Student’s t test. Representative confocal images of MCF10A-HER2 acini stained for cleaved caspase-3.

Treatment of human cancer cells with HER2 overexpression (MCF10A-HER2 or ZR-75-1) or HER2-amplified (SKBR3) (**Fig. 4d** and **Supplementary Fig. 4e**) with HC4 in 3D acini cultures in Matrigel showed that a 10-day treatment with vehicle or HC4 (2 µM) significantly increased levels of apoptosis (cleaved caspase-3) in these organoids, especially in the inner cell mass that is deprived from contact with the ECM (**Fig. 4d**). As in the *in vivo* conditions, we did not detect a significant change in the levels of proliferation as detected by phospho-histone H3 levels (**Supplementary Fig. 4f**). We conclude that early MMTV-HER2+ lesions require PERK for HER2-driven alterations in ductal epithelial organization. In HER2+ human cancer cells and mouse tumors HER2 is dependent on PERK for survival.

### PERK signaling is required for optimal HER2 phosphorylation, localization and AKT and ERK activation

We next tested the hypothesis that since HER2+ tumors are sensitive to proteotoxicity (Singh et al., 2015), inhibition of PERK might affect optimal HER2 activity due to increased ER client protein load. Detection of HER2 phosphorylation at residues Y1221/1222 in tumors showed that the area positive for P-HER2 reported by others (DiGiovanna et al., 1998) overlapped with the staining for P-PERK and P-eIF2α (**Fig. 5a**). This finding indicated that the activation of PERK and HER2 pathways co-localize. Similarly, single cell targeted-gene expression profiling of primary tumor cells also showed a population of primary tumor cells (around 25%) with high levels of ER stress genes expression (**Fig. 5b**), which could correspond to the ones showing P-HER2 activation. Importantly, when we scored the P-HER2 levels in the tumors, taking into account both the area and the intensity of the staining (**Supplementary Fig. 5a**), we found that HC4-treated tumors showed significantly lower levels of P-HER2 than control animals (**Fig. 5c**). HER2 signals as a homodimer or heterodimer with EGFR and HER3 (Moasser, 2007; Negro et al., 2004). *In vitro* treatment of MCF10A-HER2 cells that were starved and treated with EGF (100 ng/ml, 15 min) in the presence or absence of HC4 (2 µM) revealed that PERK inhibition decreased both the basal and EGF-induced levels of P-EGFR and P-HER2, along with downregulation of the survival pathway P-AKT, P-S6 and P-ERK1/2 levels (**Fig. 5d** and **Supplementary Fig. 5b**). No obvious effect was observed under these conditions on total HER2 levels or heterodimerization with EGFR as determined by surface biotinylation and co-immunoprecipitation studies (data not shown). Since HC4 does not have a direct inhibitory effect on the active site of any of the HER family members, AKT or S6 kinases (**Supplementary Table 4**), this effect must be due to an indirect effect of PERK inhibition on HER2 signaling. In contrast to other HER family members, HER2 is known to remain at the plasma membrane after ligand binding and dimerization (Hommelgaard et al., 2004; Bertelsen et al., 2014). We thus tested if HC4 might be disturbing the mechanism of activation of HER2 receptors. To this end, we performed surface biotinylation assays to measure the presence of the receptor on the cell surface, and reversible surface biotinylation to measure receptor endocytosis (Cihil et al., 2013). Our data showed that HC4 treatment decreased the amount of P-HER2 and total HER2 in the cell surface (**Fig. 5e** and **Supplementary Fig. 5c**), while concomitantly increasing endocytosed phospho-and total HER2 (**Fig. 5f**). Our data, along with previously published data (Singh et al., 2015), allow us to suggest that PERK signaling and proper UPR function is required to maintain proper HER2 downstream signaling by affecting optimal receptor localization and activation.

**Figure 5.**
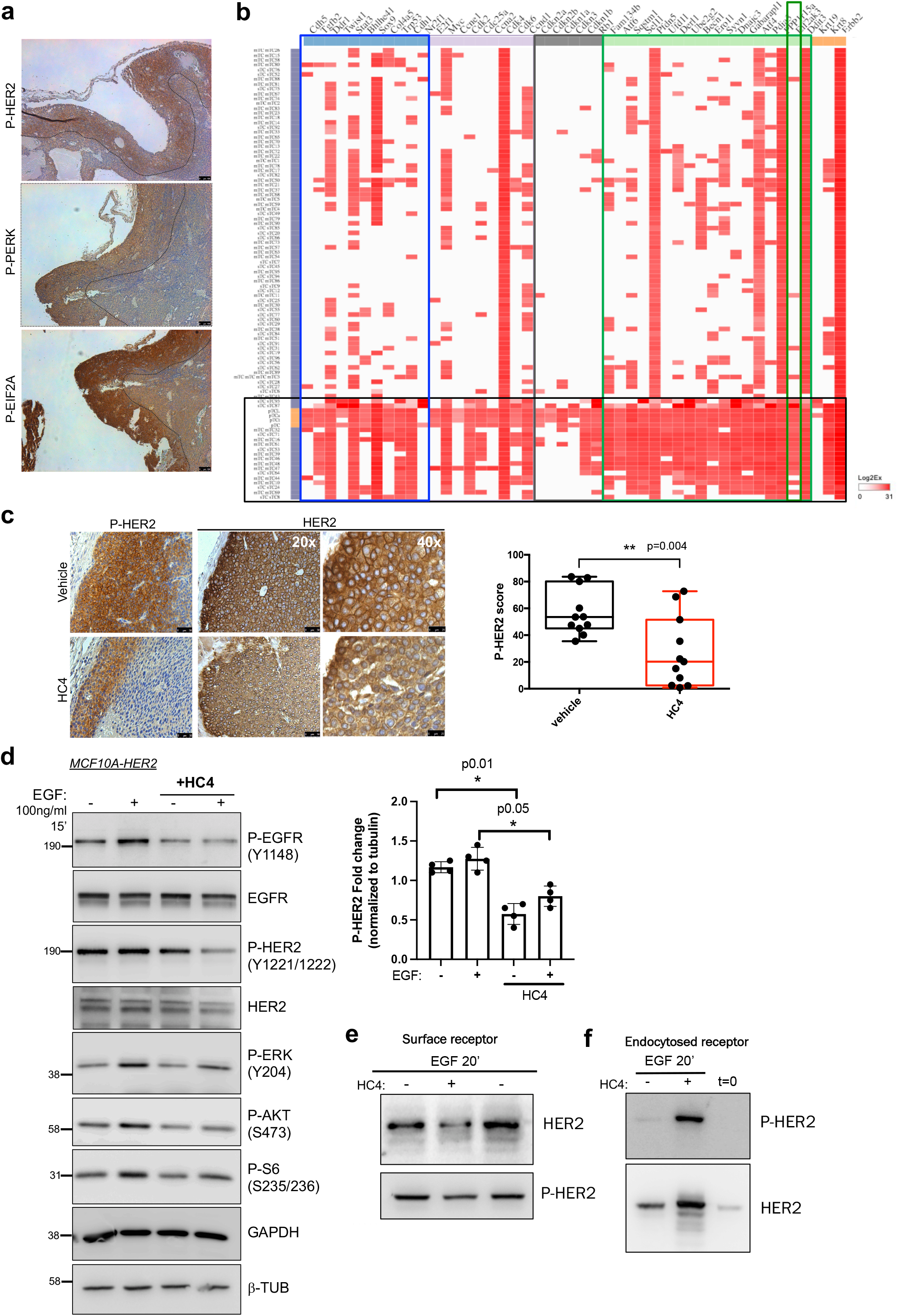
HC4 treatment decreases the levels of phospho-HER2 and downstream signaling pathways. (a) Representative images of IHC for P-HER2, P-PERK and P-EIF2α in a MMTV-HER2 breast tumor section. Note that the rim positive for P-HER2 overlaps with P-PERK and P-EIF2α stainings. Scale bar, 100 µm. (b) Hierarchical clustering of the high-throughput targeted-gene expression (columns) profile of single cells (primary breast tumor) (rows) from MMTV-HER2 females. Blue box, dormancy genes; brown box, cell cycle up genes; pink box, cell cycle down genes; green box, ER stress genes; black box, EIF2AK3 (PERK) gene. (c) Representative P-HER2 and total HER2 IHC staining in vehicle-and HC4-treated breast tumors. Quantification of P-HER2 levels in tumor sections, by IHC intensity and area scoring (N=11) (See **Supplementary Fig.5a**). Scale bar, 50 µm. P by Mann-Whitney test. (d) MCF10A-HER2 cells were starved o/n and treated +/-HC4 (2 µM), after which +/-EGF (100 ng/ml) was added for 15 min before collection. The levels of P-HER2, P-EGFR, P-AKT, P-S6 and P-ERK, as well as total HER2 and EGFR were assessed by Western blot. GAPDH and ß-tubulin were used as loading controls. Representative blot of three is shown. Densitometry analysis for P-HER2 (N=3) ± s.d. P by Student’s t test. (e) MCF10A-HER2 cells were treated as in (d) and surface receptor biotinylation assay was performed. Surface levels of total HER2 and P-HER2 were assessed. One of two experiments shown. (f) MCF10A-HER2 cells were treated as in (d) and reversible surface receptor biotinylation assay was performed. Endocytosed levels of total HER2 and P-HER2 were assessed. One of two experiments shown.

## DISCUSSION

Studies in HER2+ breast cancer models have suggested that HER2+ breast cancer tumorigenesis is dependent on PERK signaling for survival and adaptation (Bobrovnikova-Marjon et al., 2010; Singh et al., 2015). We had found that quiescent tumor cells that exist within surgical margins or as dormant DCCs in target organs (Bragado et al., 2013; Chéry et al., 2014; Sosa et al., 2014; Sosa et al., 2015) enhance their survival via PERK signaling as well as other ER stress pathways (Adomako et al., 2015; Ranganathan et al., 2008; Ranganathan et al., 2006; Schewe and Aguirre-Ghiso, 2008; Schewe and Aguirre-Ghiso, 2009). Recently, Pommier et al. validated this in their studies demonstrating that pancreatic DCCs lodged in the liver also activate a UPR during quiescence. This level of concordance across a variety of tumor types and models supports the requisite nature of this stress adaptation biology across the cancer landscape.

We now show that pharmacological PERK inhibition can selectively target HER2+ DCCs and primary lesions. A salient finding to discuss is the inhibitory effect of PERK inhibition on metastasis. In the MMTV-HER2 model, like in patients, metastases can be asynchronous with the primary tumor and sometimes develop even in instances of occult primary lesions, wherein metastases are identified earlier than the primary tumor (Husemann et al., 2008; Pavlidis and Fizazi, 2005). PERK inhibition reduced metastasis independent of the primary tumor development timeline including those initiated early (before overt tumors were palpable) as well as metastases that were coincident with overt primary tumor growth. This is important because it argues that the effect on metastasis was not simply due to reduced primary tumor burden caused by HC4. Surprisingly, metastatic burden was reduced by HC4 treatment via eliminating non-proliferative solitary or small clusters of P-Rb-negative DTCs. Imaging and single cell multiplex qPCR robustly reveal that these DCCs show an upregulation of GADD34 (protein) and a larger set of ER stress genes, including PERK itself, while also expressing genes representative of the quiescent phenotype as revealed by upregulation of several negative regulators of cell proliferation. It should be taken into account that part of the PERK-induced ER stress response is transcriptional in nature while also having a key component of preferential translation of upstream ORF-containing genes, such as ATF4 and GADD34 (Young and Wek, 2016). Similarly, UPR-induced G1 arrest has been shown to be caused by inhibiting the translation of cyclin D1 (Brewer et al., 1999). Our data strengthen the argument that quiescent DCCs are more likely to rely on PERK signaling for survival. Similarly, a sub-population of human metastatic cells from breast cancer patients also showed a negative correlation between GADD34 and Ki67, supportive of this association seen in this study in mouse models. Our data suggest that along with NR2F1 (Borgen et al.), GADD34 alone or in combination with NR2F1 may serve as a robust biomarker set for dormant/UPR^high^ DCCs and thus guide patient selection for treatment. An open question is related to the identification of the source of PERK activation in quiescent DCCs, which remains unknown.

We also demonstrate that cytostatic therapies such as CDK4/6 inhibitors (Abemaciclib) not only decrease proliferation substantially, but at the same time result in concomitant activation of the UPR as shown by high GADD34 levels in primary tumor and metastases. This observation would support the possibility of combining such CDK4/6 targeting therapies with PERK inhibitors to have an even more profound control of both proliferating and non-proliferating cancer cells. Encouragingly, the doses of HC4 we used did not significantly affect glucose levels, bone marrow or peripheral blood cell counts, drinking and feeding behavior of non-tumor or cancer bearing mice. Additional analysis revealed that HC4 treatment did not specifically alter the frequency of various innate and adaptive immune cell types (not shown), arguing that the effects we detect of HC4 on HER2 breast tumors in mice is mainly dependent on the targeting con cancer cell intrinsic pathways. Collectively these data support the dose range evaluated in which we illustrate that a significant blockade of tumor growth and metastasis is possible through the elimination of dormant DTCs and that this PERK inhibitor does not adversely affect the host’s normal organ function.

The exact mechanisms by which PERK kinase inhibition blocks tumor cell survival are unclear. It is possible that reduced adaptation to stress imposed by proteotoxicity in cancer cells (Singh et al., 2015) is a mechanism. Our data also revealed that HC4 reduced phospho-HER2 levels *in vivo* and decreased the abundance of active receptor in the membrane through enhanced endocytosis, but we did not see changes in HER2 protein degradation. However, it is still unclear how exactly PERK controls HER2 membrane localization or endocytosis. It is possible that internalization allows for better or faster de-phosphorylation of the receptor or decreases the chances of it being activated; hence resulting in decreased downstream signaling. This possibility is supported by the finding that shows that receptor endocytosis can reduce the signaling output of many plasma membrane receptors by physically reducing the concentration of the receptors at the cell surface (Sorkin et al, 2009).

In early lesions, our work also revealed that HC4 induced a differentiation phenotype. However, in established tumors, HC4 used as a single agent pushed tumors into stasis or regression via apoptosis. This argues that PERK signaling deregulation of HER2+ in early lesions is more likely linked to loss of differentiation programs, though these mechanisms have yet to be determined. Then, as the biology of the tumor progresses to become highly proliferative, the dependency on PERK signaling remains highly reliant for these HER2+ tumors.

Discovering a target and drug that can eradicate dormant DCCs is highly significant because dormant DCCs are known to evade anti-proliferative therapies *via* active and passive mechanisms (Aguirre-Ghiso et al., 2013; Naumov et al., 2003; Oshimori et al., 2015). Our work opens the door to the use of anti-dormant DTC survival therapies as a new way to target metastatic disease. This would allow targeting the full phenotypic heterogeneity of disseminated disease that may include proliferative, slow-cycling, and dormant DTCs (Aguirre-Ghiso et al., 2013). The eradication of DCCs in the bone marrow, where these cells also commonly reside in a dormant state (Bragado et al., 2013; Chéry et al., 2014; Ghajar et al., 2013; Husemann et al., 2008; Nobre et al., 2020), further strengthens the notion of PERK inhibition as an anti-dormant DCC therapy (Aguirre-Ghiso et al., 2013) that may be used in the adjuvant setting to eliminate dormant minimal residual disease (Aguirre-Ghiso et al., 2013).

## MATERIALS AND METHODS

### Reagents

EGF was obtained from PeproTech and used at 100 ng/ml. Thapsigargin was from Sigma and used at 2 nM or 0.2 µM as indicated in the legends. HC4 and Abemaciclib were provided by HiberCell and Eli Lilly, respectively.

### Cell culture

For 3D cultures, MCF10A-HER2, SKBR3 and ZR-75-1 cells were plated in growth factor-reduced Matrigel (Corning) and grown as described previously (Avivar-Valderas et al., 2013). Treatments with vehicle (DMSO) or HC4 (2 µM) were replaced every 24 h for 2D and every 48 h for 3D cultures.

### Animal work and tissue processing

Institutional Animal Care and Use Committees (IACUC) at Mount Sinai School of Medicine (MSSM) approved all animal studies. Protocol number: 08-0417. The FVB/N-Tg (MMTVneu) mouse strain was obtained from Jackson Laboratories. These mice express the un-activated neu (HER2) form under the transcriptional control of the mouse mammary tumor virus promoter/enhancer. Before being used in any experiment, females underwent one round of pregnancy and at least two weeks of no lactation after weaning. Females between 24-32 weeks of age were injected intraperitoneally with vehicle (90% corn oil, 10% ethanol) or HC4 (50 mpk) daily, for two weeks. For the combination treatment, females 24-32 weeks of age were treated daily by oral gavage with Abemaciclib (50 mpk) for 4 weeks before starting the treatment described earlier with HC4. Tumor volumes were measured using the formula (Dxd^2^)/2, where D is the longest and d is the shortest diameter. For circulating tumor cell (CTC) count, animals were anesthetized and whole blood was extracted by cardiac puncture. Mammary glands, tumors and lungs were collected and fixed in 10% buffered formalin overnight before paraffin embedding. The bone marrow from the two lower limbs was flushed with a 26 G needle and further processed by Ficoll density gradient centrifugation. For CTC as well as for disseminated tumor cell (DTC) detection in bone marrow, tissues were depleted of mature hematopoietic cells by anti-mouse antibody-labeled magnetic bead separation (Miltenyi Biotec) before fixation in formalin for 20 min at 4 °C.

### Mammary gland whole mount staining

Mammary glands fixed in 10% buffered formalin were incubated in Carmine Alum stain (Carmine 0.2%, Aluminum potassium sulfate 0.5%) (Sigma) for 2 days. Then, they were dehydrated and transfer to methyl salicylate solution before imaging using a stereomicroscope.

### Immunohistochemistry and immunofluorescence

Immunohistochemistry (IHC) and immunofluorescence (IF) from paraffin-embedded sections was performed as previously described (Avivar-Valderas et al., 2013). Briefly, slides were dewaxed and serially rehydrated. Heat-induced antigen retrieval was performed in either citrate buffer (10 mM, pH6), EDTA buffer (1 mM, pH 8) or Tris/EDTA (pH 9). Slides were further permeabilized in 0.1% Triton-X100, blocked and incubated with primary antibody overnight at 4 °C at 1:50-1:200 dilution. For IHC, an additional step of endogenous peroxidase and avidin/biotin quenching was performed before primary antibody incubation. Primary antibodies used were anti-cytokeratin 8/18 (Progen), smooth muscle actin-Cy3 (Sigma), P-PERK (T980) (provided by Eli Lilly, Tenkerian et al., 2015), P-EIF2A, Cleaved Caspase 3, P-H3 (S10) and P-HER2 (Y1221/1222) (Cell signaling), P-Rb (S249/T252) (Santa Cruz), HER2 (Abcam) and HER2 (Millipore), Ki67 (eBioscience), cytokeratin cocktail (C11 and ck7, Abcam; AE1 and AE3, Millipore) and GADD34 (Santa Cruz). Next, slides were incubated in secondary antibodies (Life Technologies) and mounted. For IHC, sections were processed using VectaStain ABC Elite kit (Vector Laboratories) and DAB Substrate kit for peroxidase labelling (Vector Laboratories) and mounted in VectaMount medium (Vector laboratories). For IF, sections were mounted in ProLong Gold Antifade aqueous medium (Thermo Fisher).

In the case of immunocytofluorescence, cytospins of fixed cells (100,00-200,000 cells/cytospin) were prepared by cyto-centrifugation at 500 rpm for 3 min on poly-prep slides, and the staining protocol was performed as explained below from the permeabilization onward.

For the staining of 3D cultures, acini were fixed in 4% PFA for 20 min at 4C, permeabilized with 0.5% Triton-X100 in PBS for 20 min at room temperature, washed in PBS-glycine and then blocked with 10% normal goat serum for 1h at 37 °C, before performing immunofluorescence staining. The scoring for P-HER2 levels is explained in Supplementary Fig.3. For the scoring of CK8/18 and SMA in mammary gland ducts, 20 low magnification fields/animal were evaluated for the expression of CK8/18 as negative (0), low (1) or high (2) and the same for SMA and the sum of the two scores was considered as the final score (from 0 to 4).

### Microscopy

Images were captured by using a Nikon Eclipse TS100 microscope, a Leica DM5500 or Leica SP5 confocal microscope.

### TUNEL *in situ* cell death detection

Apoptosis levels were evaluated using the *In situ* Cell Death Detection kit, AP (Roche). Paraffin sections from tumors were dewaxed, rehydrated and permeabilized in phosphate buffered saline (PBS) 0.2% Triton-X100 for 8 minutes. Then, slides were washed and blocked in 20% normal goat serum for 1h at 37C. The TUNEL reaction mixture was then added and let go for 1h at 37 °C. The reaction was stopped by incubating with Buffer I (0.3 M Sodium chloride, 30 mM Sodium citrate). Next, the slides were incubated with anti-fluorescein-AP antibody for 30 min. at 37 °C. After three washes in Tris buffered saline (TBS), slides were incubated in alkaline phosphatase substrate in 0.1% Tween-20 for 20 min. at room temperature. Finally, the slides were mounted using aqueous mounting medium. The percentage of TUNEL positive cells was calculated using Image J software (NIH).

### Immunoblot analysis

Cells were lysed in RIPA buffer and protein analyzed by immunoblotting as described previously (Ranganathan et al., 2006). Membranes were blotted using the additional following antibodies: P-PERK (T980) (Tenkerian et al., 2015), PERK (Santa Cruz), P-EGFR (Y1148), EGFR, P-AKT (S473), P-S6 (S235/236), P-ERK (Y204), P-HER2 (Y1221/1222, Y1112, Y877), HSP90 (Cell signaling), GAPDH (Millipore) and β-Tubulin (Abcam). For induction of ER stress, MCF10A-HER2 cells were plated in low adhesion plates for 24h before collection.

### Cell surface biotinylation and endocytosis assay

For cell surface biotinylation, we used Pierce cell surface protein isolation kit following manufacturer’s instructions with minor changes. Briefly, MCF10A-HER2 cells were serum- and EGF-starved and treated +/- HC4 for 24h before being stimulated with +/- EGF (100 ng/ml) for 20’. Then, cells were washed with ice-cold PBS and surface proteins biotinylated for 30 min at 4C. After quenching, cells were harvested and lysed using RIPA buffer. Protein lysates were incubated with NeutrAvidin agarose beads and the bound proteins were released by incubation with SDS-PAGE sample buffer containing DTT (50 mM). For endocytosis assays (Cihil et al., 2013), cells were treated similarly but before treatment with EGF cell surface proteins were biotinylated. After 20 min incubation +/- EGF (100 ng/ml) at 37C (to induce endocytosis), cells were washed with ice-cold PBS and incubated with stripping buffer (to remove cell surface biotinylation: 75 mM NaCl, 1mM MgCl2, 0.1mM CaCl2, 50 mM glutathione and 80 mM NaOH, pH 8.6) for 30’. To control for stripping efficiency, cells were stripped without 37C incubation (t=0). Cell lysates were prepared and processed for biotinylated protein isolation as described before.

### Single cell targeted gene expression analysis

Primary tumors from MMTV-neu 28-30-week old females were digested with collagenase into a single cell suspension. Lungs from MMTV-neu 24-30-week old females were digested into a single cell suspension with collagenase and resuspended in FACS buffer. Cells were then stained with anti-HER2-PE, anti-CD45-APC and DAPI and the HER2+/CD45-population of cells sorted using a BDFACSAria sorter as previously described (Aguirre-Ghiso et al., 2021). Sorted cells were resuspended at a 312,500cells/ml concentration in media and 80 ul were mixed with 20 ul suspension reagent (C1 Fluidigm). A C1 Single-cell Preamp IFC 10-17 um was used for the single cell separation. Pre-amplification was run using Ambion Single Cell-to-CT qRT-PCR kit and 20x TaqMan Gene expression FAM-MGB assays. Resulting cDNA was further diluted in C1 DNA dilution reagent 1/3 and used for gene expression analysis using 96.96 IFCs (Fluidigm), Juno System controller and Biomark HD for high-throughput qPCR. TaqMan Fast Advanced Master Mix was used for the qPCR reactions. Analysis was performed using Fluidigm Real-Time PCR Analysis Software and Clustergrammer web-based tool (Fernandez *et al*., 2017) for hierarchical clustering heatmaps.

### Database

TCGA data on mRNA expression levels of EIF2AK3 was accessed and analyzed through cBioPortal (https://bit.ly/3yBhw2b).

### Statistical analysis

All points represent independent biological samples with error bars representing standard deviations and statistical significance was determined using one-sided Mann–Whitney test using the Graph Pad Prism Software.

## ACKNOWLEDGEMENTS

We thank the Aguirre-Ghiso and HiberCell teams for useful discussions. We thank Brian Lee for help with Clustergrammer use. Grant Support: Eli Lilly to J.A.A-G, LIFA Fellowship (Eli Lilly) to V.C., NIH/NCI (CA109182, CA196521, CA216248), to J.A.A-G, HiberCell and DoD-BCRP Breakthrough Award (BC132674) to J.A.A-G. JA.A-G is a Samuel Waxman Cancer Research Foundation Investigator.

## AUTHOR CONTRIBUTIONS

VC and WZ designed, planned and conducted experiments, analyzed data, and wrote the manuscript; VC, EFF, WZ, ARN, JC performed *in vivo* mouse experiments. WZ and VC performed immune profiling experiments and pharmacokinetic studies. KS, AN, MM and ACR designed, developed and directed all pharmacology related to PERK inhibitors and participated in experimental design and analyzed data; JAAG conceived the project and designed experiments. VC, WZ, MM, ACR and JAAG analyzed data, provided insight, wrote and edited the manuscript.

## REFERENCES

Adomako, A., Calvo, V., Biran, N., Osman, K., Chari, A., Paton, J. C., Paton, A. W., Moore, K., Schewe, D. M., and Aguirre-Ghiso, J. A. (2015). Identification of markers that functionally define a quiescent multiple myeloma cell sub-population surviving bortezomib treatment. BMC Cancer 15, 444.

Aguirre-Ghiso, J. A., Bragado, P., and Sosa, M. S. (2013). Metastasis awakening: targeting dormant cancer. Nat Med 19, 276–277.

Avivar-Valderas, A., Bobrovnikova-Marjon, E., Alan Diehl, J., Bardeesy, N., Debnath, J., and Aguirre-Ghiso, J. A. (2013). Regulation of autophagy during ECM detachment is linked to a selective inhibition of mTORC1 by PERK. Oncogene.

Avivar-Valderas, A., Salas, E., Bobrovnikova-Marjon, E., Diehl, J. A., Nagi, C., Debnath, J., and Aguirre-Ghiso, J. A. (2011). PERK integrates autophagy and oxidative stress responses to promote survival during extracellular matrix detachment. Mol Cell Biol 31, 3616–3629.

Bi, M., Naczki, C., Koritzinsky, M., Fels, D., Blais, J., Hu, N., Harding, H., Novoa, I., Varia, M., Raleigh, J., et al. (2005). ER stress-regulated translation increases tolerance to extreme hypoxia and promotes tumor growth. Embo J 24, 3470–3481.

Blais, J. D., Filipenko, V., Bi, M., Harding, H. P., Ron, D., Koumenis, C., Wouters, B. G., and Bell, J. C. (2004). Activating transcription factor 4 is translationally regulated by hypoxic stress. Mol Cell Biol 24, 7469–7482.

Bobrovnikova-Marjon, E., Grigoriadou, C., Pytel, D., Zhang, F., Ye, J., Koumenis, C., Cavener, D., and Diehl, J. A. PERK promotes cancer cell proliferation and tumor growth by limiting oxidative DNA damage. Oncogene 29, 3881–3895.

Bobrovnikova-Marjon, E., Grigoriadou, C., Pytel, D., Zhang, F., Ye, J., Koumenis, C., Cavener, D., and Diehl, J. A. (2010). PERK promotes cancer cell proliferation and tumor growth by limiting oxidative DNA damage. Oncogene 29, 3881–3895.

Bobrovnikova-Marjon, E., Hatzivassiliou, G., Grigoriadou, C., Romero, M., Cavener, D. R., Thompson, C. B., and Diehl, J. A. (2008). PERK-dependent regulation of lipogenesis during mouse mammary gland development and adipocyte differentiation. Proc Natl Acad Sci U S A 105, 16314–16319.

Bragado, P., Estrada, Y., Parikh, F., Krause, S., Capobianco, C., Farina, H. G., Schewe, D. M., and Aguirre-Ghiso, J. A. (2013). TGF-beta2 dictates disseminated tumour cell fate in target organs through TGF-beta-RIII and p38alpha/beta signalling. Nat Cell Biol 15, 1351–1361.

Cerami, E., Gao, J., Dogrusoz, U., Gross, B. E., Sumer, S. O., Aksoy, B. A., Jacobsen, A., Byrne, C. J., Heuer, M. L., Larsson, E., et al. (2012). The cBio cancer genomics portal: an open platform for exploring multidimensional cancer genomics data. Cancer discovery 2, 401–404.

Chen, X., Iliopoulos, D., Zhang, Q., Tang, Q., Greenblatt, M. B., Hatziapostolou, M., Lim, E., Tam, W. L., Ni, M., Chen, Y., et al. (2014). XBP1 promotes triple-negative breast cancer by controlling the HIF1alpha pathway. Nature 508, 103–107.

Chéry, L., Lam, H.-M., Coleman, I., Lakely, B., Coleman, R., Larson, S., Aguirre-Ghiso, J. A., Xia, J., Gulati, R., Nelson, P. S., et al. (2014). Characterization of single disseminated prostate cancer cells reveals tumor cell heterogeneity and identifies dormancy associated pathways).

Chevet, E., Hetz, C., and Samali, A. (2015). Endoplasmic reticulum stress-activated cell reprogramming in oncogenesis. Cancer Discov 5, 586–597.

DiGiovanna, M. P., Lerman, M. A., Coffey, R. J., Muller, W. J., Cardiff, R. D., and Stern, D. F. (1998). Active signaling by Neu in transgenic mice. Oncogene 17, 1877–1884.

Espina, V., Mariani, B. D., Gallagher, R. I., Tran, K., Banks, S., Wiedemann, J., Huryk, H., Mueller, C., Adamo, L., Deng, J., et al. (2010). Malignant precursor cells pre-exist in human breast DCIS and require autophagy for survival. PloS one 5, e10240.

Ghajar, C. M., Peinado, H., Mori, H., Matei, I. R., Evason, K. J., Brazier, H., Almeida, D., Koller, A., Hajjar, K. A., Stainier, D. Y., et al. (2013). The perivascular niche regulates breast tumour dormancy. Nat Cell Biol 15, 807–817.

Guy, C. T., Webster, M. A., Schaller, M., Parsons, T. J., Cardiff, R. D., and Muller, W. J. (1992). Expression of the neu protooncogene in the mammary epithelium of transgenic mice induces metastatic disease. Proc Natl Acad Sci U S A 89, 10578–10582.

Harper, K. L., Sosa, M. S., Entenberg, D., Hosseini, H., Cheung, J. F., Nobre, R., Avivar-Valderas, A., Nagi, C., Girnius, N., Davis, R. J., et al. (2016). Mechanism of early dissemination and metastasis in Her2(+) mammary cancer. Nature 540, 588–592.

Hart, L. S., Cunningham, J. T., Datta, T., Dey, S., Tameire, F., Lehman, S. L., Qiu, B., Zhang, H., Cerniglia, G., Bi, M., et al. (2012). ER stress-mediated autophagy promotes Myc-dependent transformation and tumor growth. J Clin Invest 122, 4621–4634.

Husemann, Y., Geigl, J. B., Schubert, F., Musiani, P., Meyer, M., Burghart, E., Forni, G., Eils, R., Fehm, T., Riethmuller, G., and Klein, C. A. (2008). Systemic spread is an early step in breast cancer. Cancer Cell 13, 58–68.

Lu, Y., Bertran, S., Samuels, T. A., Mira-y-Lopez, R., and Farias, E. F. (2010). Mechanism of inhibition of MMTV-neu and MMTV-wnt1 induced mammary oncogenesis by RARalpha agonist AM580. Oncogene 29, 3665–3676.

Moasser, M. M. (2007). The oncogene HER2: its signaling and transforming functions and its role in human cancer pathogenesis. Oncogene 26, 6469–6487.

Muller, W. J., Sinn, E., Pattengale, P. K., Wallace, R., and Leder, P. (1988). Single-step induction of mammary adenocarcinoma in transgenic mice bearing the activated c-neu oncogene. Cell 54, 105–115.

Naumov, G. N., Townson, J. L., MacDonald, I. C., Wilson, S. M., Bramwell, V. H., Groom, A. C., and Chambers, A. F. (2003). Ineffectiveness of doxorubicin treatment on solitary dormant mammary carcinoma cells or late-developing metastases. Breast Cancer Res Treat 82, 199–206.

Negro, A., Brar, B. K., and Lee, K. F. (2004). Essential roles of Her2/erbB2 in cardiac development and function. Recent progress in hormone research 59, 1–12.

Nobre, A.R., Dalla, E., Yang, J., Huang, X., Kenigsberg, E., Wang, J. and Aguirre-Ghiso, J.(2021). A Mesenchymal-like Program of Dormancy controlled by ZFP281 Serves as a Barrier To Metastatic Progression of Early Disseminated Cancer Cells.

Nobre, A. R., Risson, E., Singh, D. K., Di Martino, J., Cheung, J. F., Wang, J., Johnson, J., Russnes, H. G., Bravo-Cordero, J. J., Birbrair, A., et al. (2020). NG2+/Nestin+ mesenchymal stem cells dictate DTC dormancy in the bone marrow through TGFβ2. bioRxiv Online, 2020.2010.2022.349514.

Oshimori, N., Oristian, D., and Fuchs, E. (2015). TGF-beta promotes heterogeneity and drug resistance in squamous cell carcinoma. Cell 160, 963–976.

Ozcan, U., Ozcan, L., Yilmaz, E., Duvel, K., Sahin, M., Manning, B. D., and Hotamisligil, G. S. (2008). Loss of the tuberous sclerosis complex tumor suppressors triggers the unfolded protein response to regulate insulin signaling and apoptosis. Mol Cell 29, 541–551.

Pavlidis, N., and Fizazi, K. (2005). Cancer of unknown primary (CUP). Crit Rev Oncol Hematol 54, 243–250.

Ranganathan, A. C., Ojha, S., Kourtidis, A., Conklin, D. S., and Aguirre-Ghiso, J. A. (2008). Dual function of pancreatic endoplasmic reticulum kinase in tumor cell growth arrest and survival. Cancer Res 68, 3260–3268.

Ranganathan, A. C., Zhang, L., Adam, A. P., and Aguirre-Ghiso, J. A. (2006). Functional coupling of p38-induced up-regulation of BiP and activation of RNA-dependent protein kinase-like endoplasmic reticulum kinase to drug resistance of dormant carcinoma cells. Cancer Res 66, 1702–1711.

Romero-Ramirez, L., Cao, H., Regalado, M. P., Kambham, N., Siemann, D., Kim, J. J., Le, Q. T., and Koong, A. C. (2009). X box-binding protein 1 regulates angiogenesis in human pancreatic adenocarcinomas. Transl Oncol 2, 31–38.

Rouschop, K. M., van den Beucken, T., Dubois, L., Niessen, H., Bussink, J., Savelkouls, K., Keulers, T., Mujcic, H., Landuyt, W., Voncken, J. W., et al. (2010). The unfolded protein response protects human tumor cells during hypoxia through regulation of the autophagy genes MAP1LC3B and ATG5. J Clin Invest 120, 127–141.

Schewe, D. M., and Aguirre-Ghiso, J. A. (2008). ATF6alpha-Rheb-mTOR signaling promotes survival of dormant tumor cells in vivo. Proc Natl Acad Sci U S A 105, 10519–10524.

Schewe, D. M., and Aguirre-Ghiso, J. A. (2009). Inhibition of eIF2alpha dephosphorylation maximizes bortezomib efficiency and eliminates quiescent multiple myeloma cells surviving proteasome inhibitor therapy. Cancer Res 69, 1545–1552.

Singh, N., Joshi, R., and Komurov, K. (2015). HER2-mTOR signaling-driven breast cancer cells require ER-associated degradation to survive. Sci Signal 8, ra52.

Sosa, M. S., Bragado, P., and Aguirre-Ghiso, J. A. (2014). Mechanisms of disseminated cancer cell dormancy: an awakening field. Nat Rev Cancer 14, 611–622.

Sosa, M. S., Parikh, F., Maia, A. G., Estrada, Y., Bosch, A., Bragado, P., Ekpin, E., George, A., Zheng, Y., Lam, H. M., et al. (2015). NR2F1 controls tumour cell dormancy via SOX9-and RARbeta-driven quiescence programmes. Nat Commun 6, 6170.

Tameire, F., Verginadis, II, and Koumenis, C. (2015). Cell intrinsic and extrinsic activators of the unfolded protein response in cancer: Mechanisms and targets for therapy. Semin Cancer Biol 33, 3–15.

Tenkerian, C., Krishnamoorthy, J., Mounir, Z., Kazimierczak, U., Khoutorsky, A., Staschke, K. A., Kristof, A. S., Wang, S., Hatzoglou, M., and Koromilas, A. E. (2015). mTORC2 Balances AKT Activation and eIF2alpha Serine 51 Phosphorylation to Promote Survival under Stress. Mol Cancer Res 13, 1377–1388.

Walter, P., and Ron, D. (2011). The unfolded protein response: from stress pathway to homeostatic regulation. Science 334, 1081–1086.

Ye, J., Kumanova, M., Hart, L. S., Sloane, K., Zhang, H., De Panis, D. N., Bobrovnikova-Marjon, E., Diehl, J. A., Ron, D., and Koumenis, C. The GCN2-ATF4 pathway is critical for tumour cell survival and proliferation in response to nutrient deprivation. Embo J 29, 2082–2096.

Yu, Q., Zhao, B., Gui, J., Katlinski, K. V., Brice, A., Gao, Y., Li, C., Kushner, J. A., Koumenis, C., Diehl, J. A., and Fuchs, S. Y. (2015). Type I interferons mediate pancreatic toxicities of PERK inhibition. Proc Natl Acad Sci U S A 112, 15420–15425.

