## Supplemental figures and legends for "PERK inhibition blocks metastasis initiation by limiting UPR-dependent survival of dormant disseminated cancer cells"

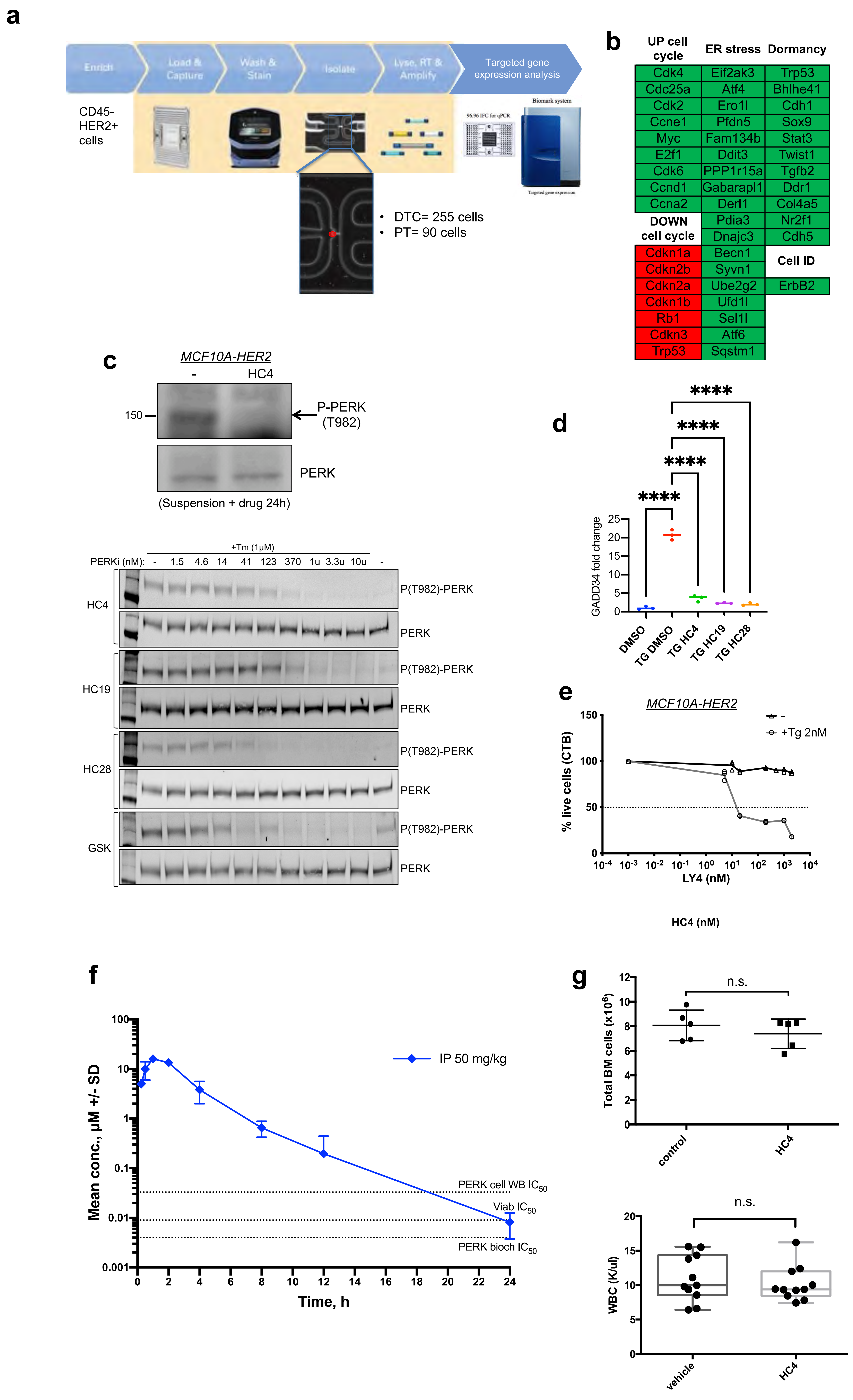


**Supplementary Figure 1**

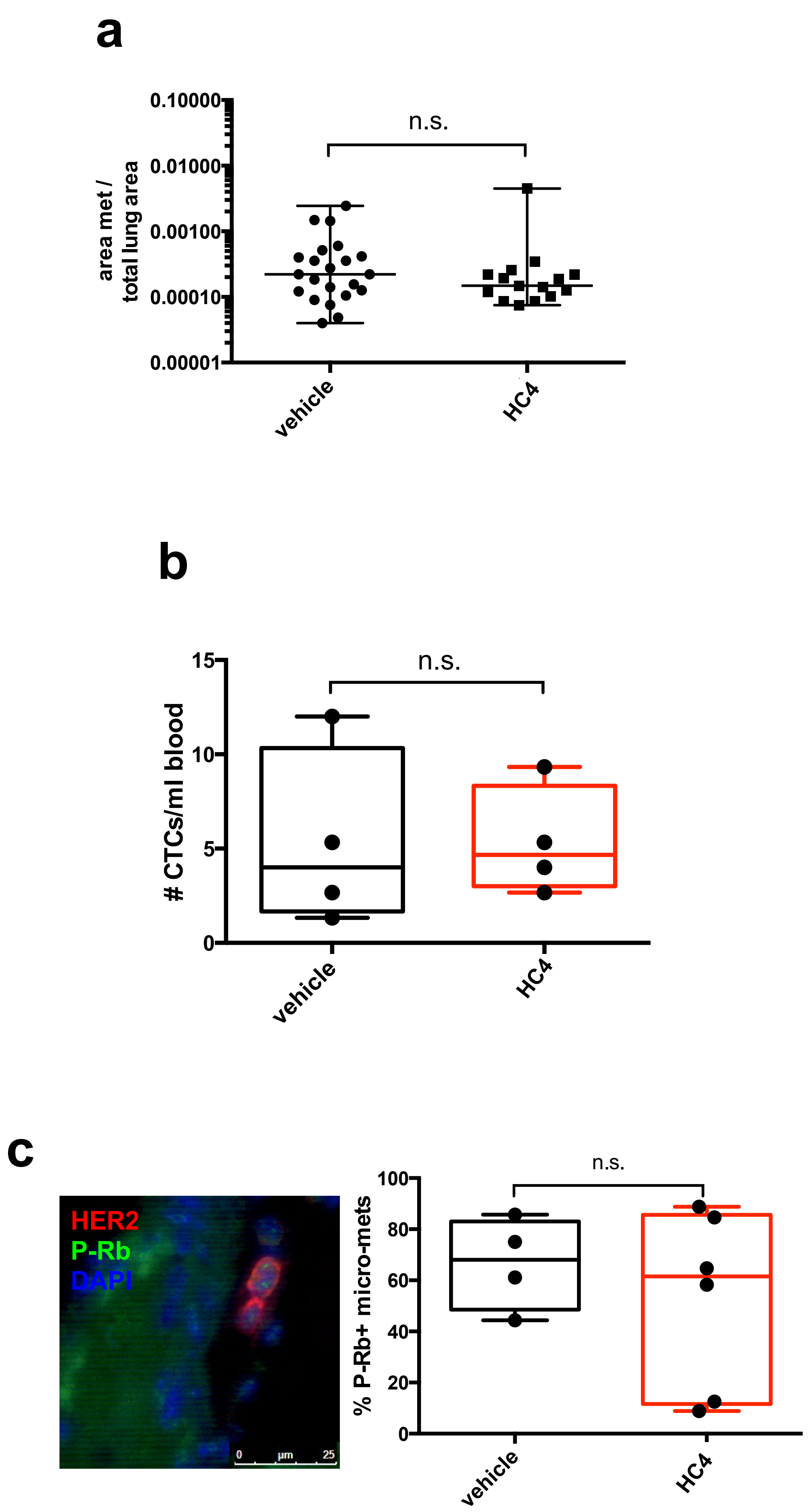


**Supplementary Figure 2**

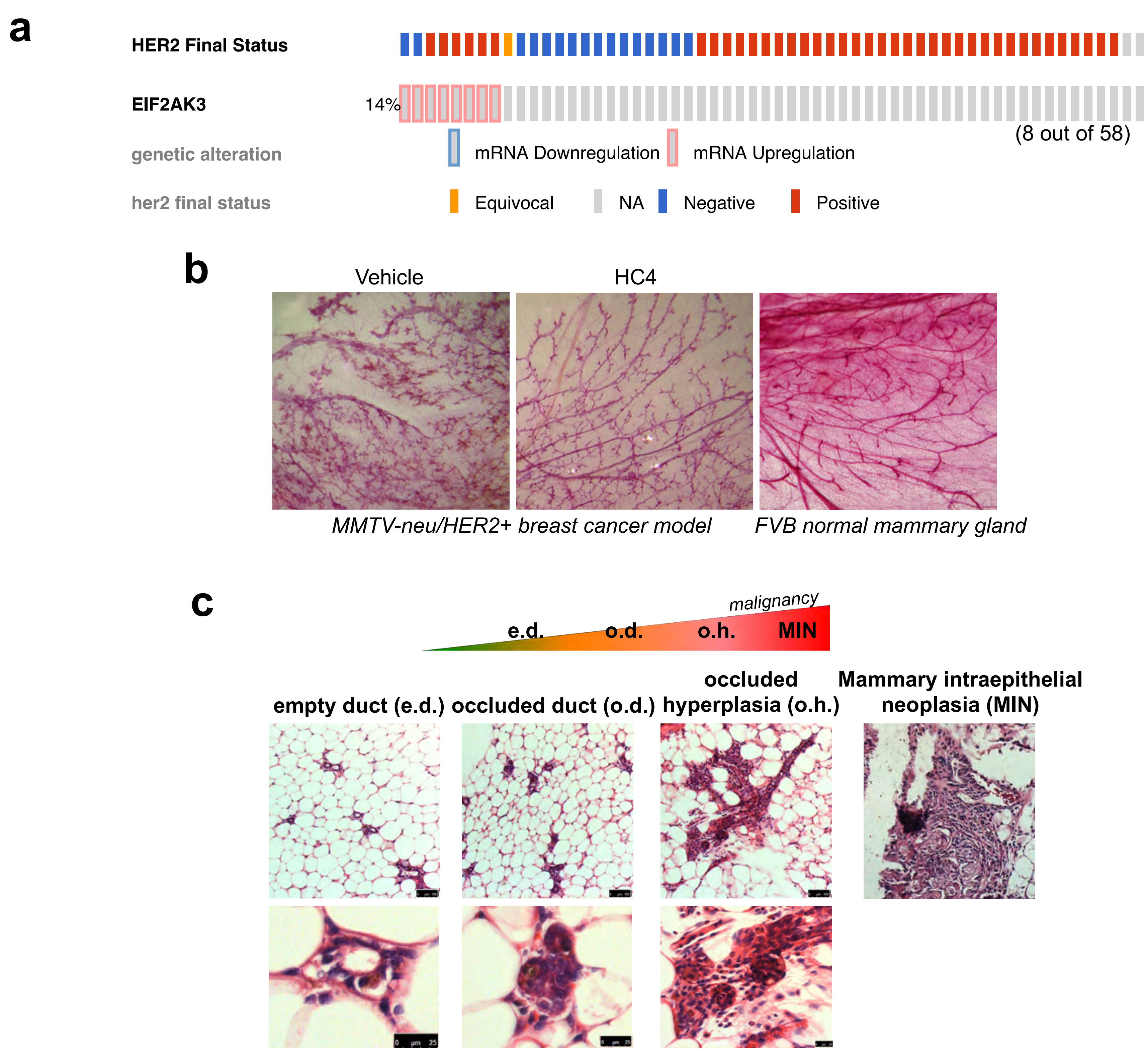


**Supplementary Figure 3**

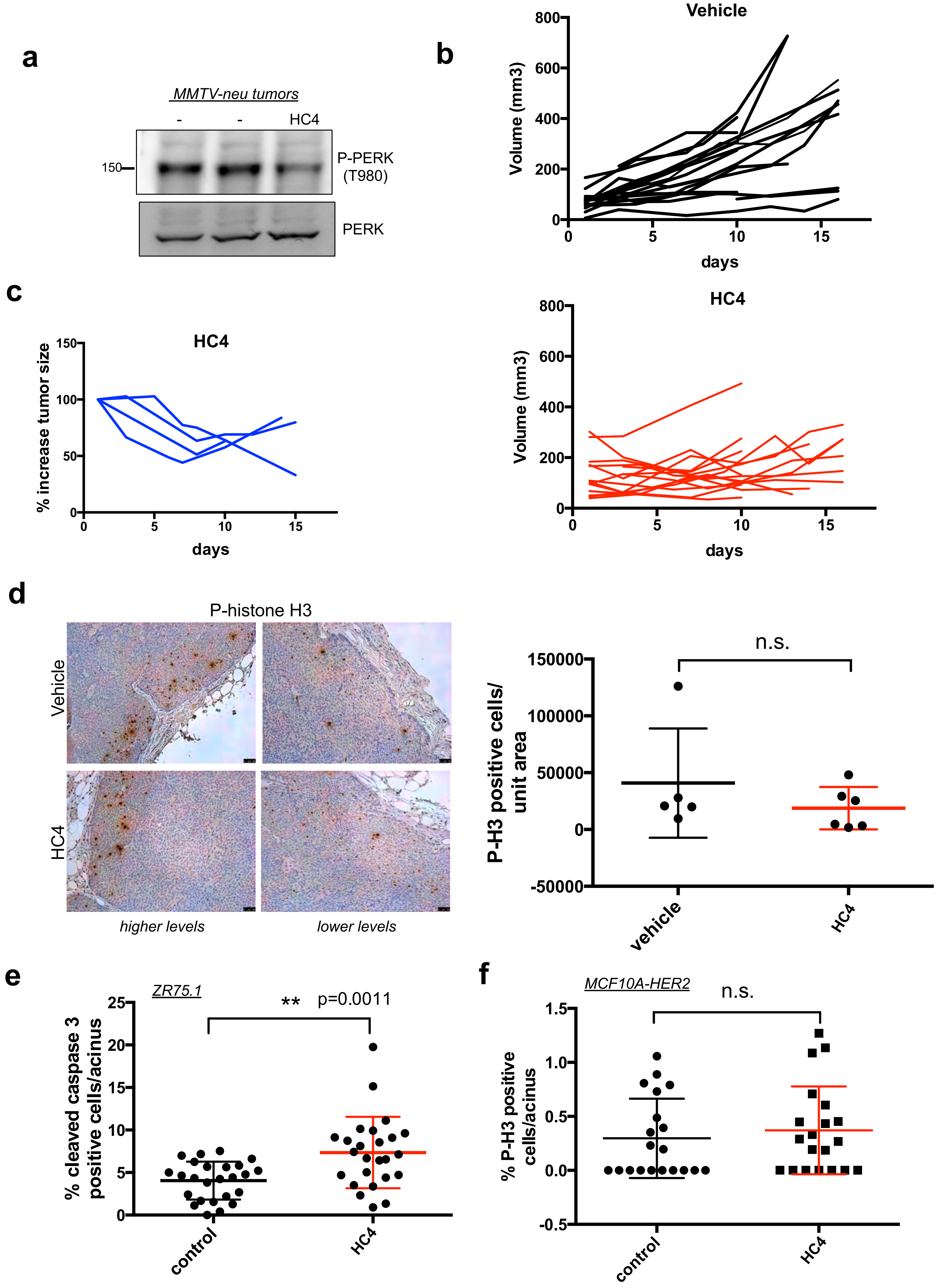


**Supplementary Figure 4**

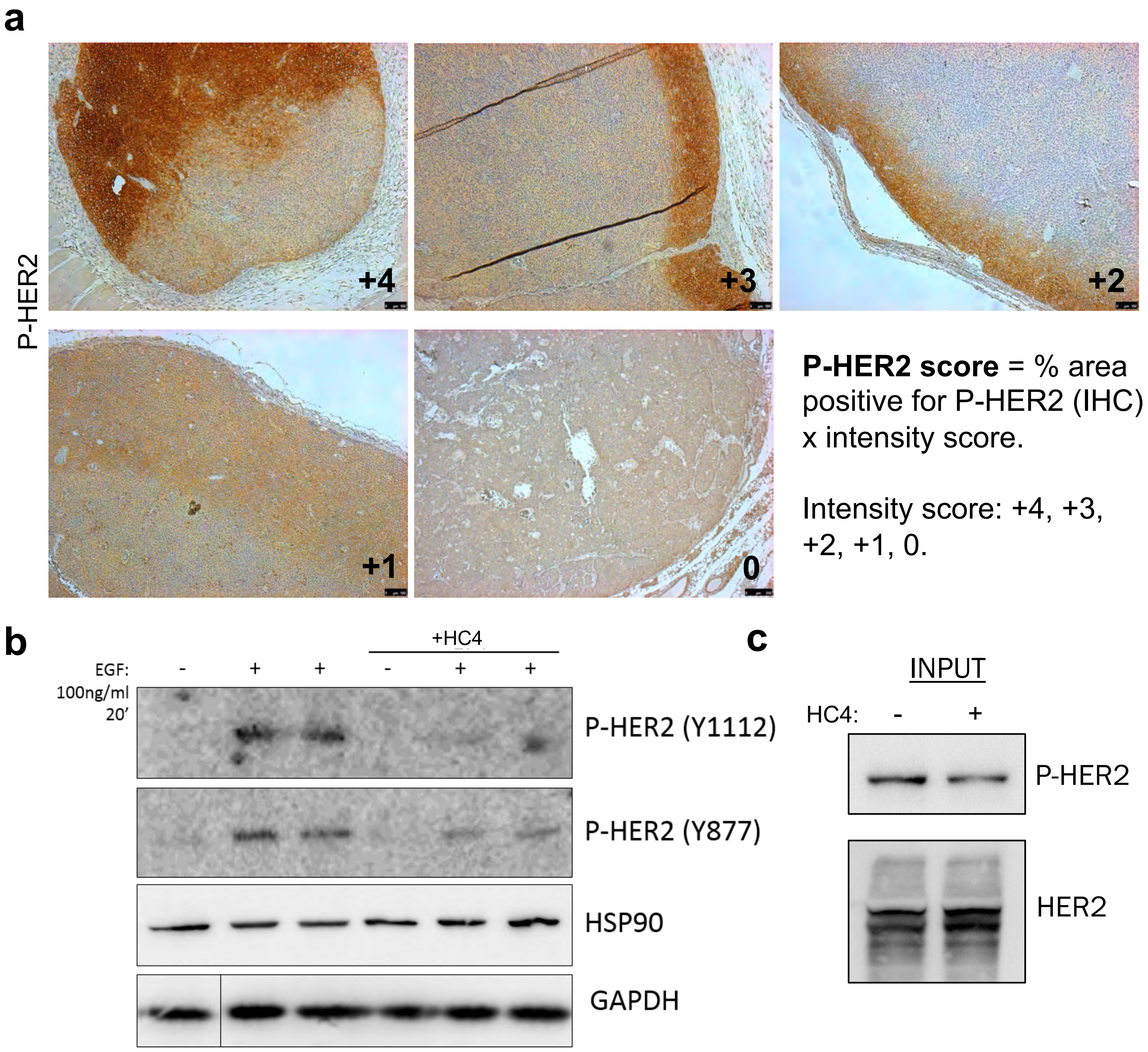


**Supplementary Figure 5**

**Supplementary Table 1 Human breast cancer metastases samples**

| Patient | Name | ER status | PR status | Her2 status | Pathology | Metastasis sample | Cytokeratins | Ki67 | GADD34 |
|---|---|---|---|---|---|---|---|---|---|
| 5 | 14850 | Neg | Neg | Pos | Ductal carcinoma | lymph node | positive | high | low |
| 4 | 14851 | Pos | Neg | Pos | Ductal carcinoma | lymph node | positive | low | intermediate |
| 7 | 14852 | Neg | Neg | Pos | Ductal carcinoma | lymph node | positive | high | low |
| 8 | 14853 | Neg | Neg | Pos | Ductal carcinoma | lymph node | positive | high | low |
| 6 | 14854 | Pos | Pos | Pos | Ductal carcinoma | lymph node | positive | intermediate | low |
| 1 | 14855 | Pos | Pos | Pos | Ductal carcinoma | lymph node | positive | low | high |
| 3 | 14856 | Pos | Pos | Pos | Ductal carcinoma | lymph node | positive | low | high |
| 9 | 14857 | Pos | Pos | Pos | Ductal carcinoma | lymph node | positive | high | negative |
| 10 | 14858 | Neg | Neg | Pos | Ductal carcinoma | lymph node | positive | intermediate | negative |
| 2 | 14859 | Pos | Neg | Pos | Ductal carcinoma | lymph node | positive | low | high |
| | 9876 part D | Pos | Pos | Neg | Invasive poorly differentiated | lymph node | positive | low | intermediate |
| | 3603 part B | Pos | Pos | Neg | Ductal moderately differentiated | liver | positive | high | low |
| | 879 part C | Neg | Neg | Neg | Not stated | liver | positive | high | negative |
| | 1645 part F | Neg | Neg | Pos | Ductal moderately differentiated | liver | positive | low | high |
| | 1415 part B | Pos | Pos | Neg | Ductal moderately differentiated | liver | positive | intermediate | intermediate |
| | 8852 part A | N/A | N/A | N/A | Malignant phyllodes, high grade sarcomatoid | chest wall | negative | low | low |
| | 943 part C | Pos, weak | Neg | Neg | Adeno poorly differentiated | lung | positive | low | high |

**Supplementary Table 2: Enzymatic and cell based $IC_{50}$ values related to PERK, ATF4 and GCN2**

| Compound | PERK[a] Enzyme $IC_{50}$ (μM) | PERK[b] Cell-based $IC_{50}$ (μM) | PERK-activated ATF4-luc[c] Cell-based $IC_{50}$ (μM) | GCN2-activated ATF4-luc[d] Cell-based $IC_{50}$ (μM) |
|---|---|---|---|---|
| HC4 | 0.004 | 0.033 | 0.067 | >10 |
| HC19 | 0.004 | 0.045 | 0.072 | >10 |
| HC28 | 0.002 | 0.009 | 0.026 | >10 |
| GSK157 | 0.002 | 0.014 | 0.024 | >10 |

[a]PERK biochemical assay using purified eIF2a as substrate. [b]Cell-based assay of tunicamycin-induced eIF2a phosphorylation in 293 cells. [c]Cell-based assay of tunicamycin-induced (PERK activated) ATF4-Luc activity in 293 cells. [d]GCN2 Cell-based assay of Halofuginone-induced (GCN2 activated) ATF4-Luc activity in 293 cells.

**Supplementary Table 3: KINOMEscan™ screening comparison of HC4, HC19, HC28 and GSK-157 at concentrations of 0.1, 1.0 and 10 μM versus other eif2α kinases**

| KINOMEscan Gene Symbol | Entrez Gene Symbol | GSK-157 | | | HC4 | | | HC19 | | | HC28 | | |
|---|---|---|---|---|---|---|---|---|---|---|---|---|---|
| | | 0.1 μM | 1.0 μM | 10.0 μM | 0.1 μM | 1.0 μM | 10.0 μM | 0.1 μM | 1.0 μM | 10.0 μM | 0.1 μM | 1.0 μM | 10.0 μM |
| EIF2AK1 | EIF2AK1 | 85 | 25 | 29 | 93 | 100 | 40 | 100 | 100 | 80 | 100 | 95 | 28 |
| PRKR | EIF2AK2 | 81 | 72 | 29 | 100 | 100 | 100 | 100 | 100 | 100 | 100 | 100 | 100 |
| GCN2(Kin.Dom.2,S808G) | EIF2AK4 | 96 | 73 | 26 | 88 | 90 | 64 | 87 | 78 | 52 | 77 | 69 | 11 |

The compounds were screened at concentrations of 0.1, 1.0 and10 mM , and results for primary screen binding interactions are reported as '% Ctrl', where lower numbers indicate stronger hits in the matrix. Red highlights those hits as <30% of Ctrl.

**Supplementary Table 4: KINOMEscan™ screening comparison of HC4, HC19, HC28 and GSK-157 at concentrations of 0.1, 1.0 and 10 μM versus select kinases in the HER family signaling pathway**

| KINOMEscan Gene Symbol | Entrez Gene Symbol | GSK-157 | | | HC4 | | | HC19 | | | HC28 | | |
|---|---|---|---|---|---|---|---|---|---|---|---|---|---|
| | | 0.1 μM | 1.0 μM | 10.0 μM | 0.1 μM | 1.0 μM | 10.0 μM | 0.1 μM | 1.0 μM | 10.0 μM | 0.1 μM | 1.0 μM | 10.0 μM |
| AKT1 | AKT1 | 100 | 96 | 87 | 93 | 91 | 95 | 100 | 100 | 100 | 100 | 100 | 100 |
| AKT2 | AKT2 | 94 | 100 | 95 | 94 | 100 | 100 | 100 | 96 | 98 | 100 | 100 | 100 |
| AKT3 | AKT3 | 100 | 81 | 82 | 97 | 95 | 87 | 100 | 100 | 96 | 100 | 100 | 100 |
| EGFR | EGFR | 100 | 100 | 92 | 100 | 79 | 77 | 100 | 99 | 100 | 100 | 100 | 98 |
| ERBB2 | ERBB2 | 91 | 85 | 24 | 94 | 100 | 65 | 85 | 88 | 75 | 79 | 81 | 80 |
| ERBB3 | ERBB3 | 88 | 96 | 87 | 92 | 91 | 90 | 99 | 100 | 92 | 83 | 93 | 89 |
| ERBB4 | ERBB4 | 68 | 77 | 74 | 90 | 83 | 91 | 100 | 100 | 100 | 98 | 100 | 100 |
| ERK1 | MAPK3 | 90 | 100 | 100 | 96 | 96 | 99 | 96 | 93 | 99 | 100 | 100 | 100 |
| ERK2 | MAPK1 | 99 | 98 | 96 | 100 | 91 | 96 | 100 | 91 | 100 | 100 | 100 | 100 |
| ERK3 | MAPK6 | 91 | 100 | 100 | 85 | 100 | 84 | 100 | 100 | 100 | 100 | 100 | 99 |
| ERK4 | MAPK4 | 100 | 100 | 96 | 89 | 98 | 91 | 97 | 100 | 100 | 100 | 100 | 100 |
| ERK5 | MAPK7 | 100 | 91 | 100 | 98 | 95 | 91 | 100 | 100 | 100 | 100 | 100 | 100 |
| ERK8 | MAPK15 | 98 | 66 | 81 | 100 | 82 | 88 | 99 | 100 | 100 | 100 | 80 | 84 |
| S6K1 | RPS6KB1 | 80 | 83 | 50 | 84 | 89 | 82 | 93 | 91 | 88 | 100 | 99 | 100 |
| SRC | SRC | 88 | 60 | 7 | 93 | 95 | 86 | 100 | 100 | 100 | 100 | 100 | 97 |
| SYK | SYK | 97 | 76 | 92 | 81 | 65 | 84 | 100 | 100 | 100 | 100 | 100 | 82 |
| CSK | CSK | 89 | 97 | 41 | 95 | 96 | 81 | 98 | 94 | 89 | 97 | 98 | 93 |
| PIK3CA | PIK3CA | 98 | 90 | 91 | 95 | 89 | 100 | 100 | 100 | 100 | 100 | 100 | 100 |
| PIK3CB | PIK3CB | 97 | 100 | 100 | 100 | 100 | 97 | 84 | 91 | 70 | 91 | 90 | 89 |
| PIK3CG | PIK3CG | 92 | 76 | 72 | 76 | 88 | 60 | 100 | 100 | 75 | 100 | 100 | 100 |
| PIK3C2B | PIK3C2B | 63 | 94 | 5 | 85 | 73 | 7 | 100 | 40 | 1 | 100 | 59 | 25 |
| PYK2 | PTK2B | 91 | 84 | 85 | 91 | 89 | 93 | 100 | 100 | 100 | 100 | 100 | 100 |

The compounds were screened at concentrations of 0.1, 1.0 and10 mM , and results for primary screen binding interactions are reported as '% Ctrl', where lower numbers indicate stronger hits in the matrix. Red highlights those hits as <30% of Ctrl.

**Supplementary Table 5: KINOMEscan™ screening comparison of HC4, HC19, HC28 and GSK-157 at concentrations of 0.1, 1.0 and 10 mM versus RIPK1, AURKB, AXL, EPHB6, FLT3, KIT and KIT (V599D), kinases potently engaged by GSK157**

| KINOMEscan Gene Symbol | Entrez Gene Symbol | GSK-157 | | | HC4 | | | HC19 | | | HC28 | | |
|---|---|---|---|---|---|---|---|---|---|---|---|---|---|
| | | 0.1 μM | 1.0 μM | 10.0 μM | 0.1 μM | 1.0 μM | 10.0 μM | 0.1 μM | 1.0 μM | 10.0 μM | 0.1 μM | 1.0 μM | 10.0 μM |
| RIPK1 | | 15 | 0.1 | 0 | 97 | 100 | 100 | 100 | 100 | 98 | 100 | 100 | 100 |
| AURKB | | 80 | 34 | 5 | 78 | 80 | 82 | 97 | 96 | 81 | 78 | 87 | 75 |
| AXL | | 41 | 19 | 8 | 94 | 95 | 70 | 93 | 95 | 77 | 100 | 94 | 70 |
| EPHB6 | | 83 | 41 | 5 | 98 | 100 | 87 | 100 | 100 | 96 | 100 | 100 | 79 |
| FLT3 | | 90 | 63 | 11 | 100 | 99 | 66 | 96 | 91 | 73 | 98 | 100 | 55 |
| KIT | | 49 | 6 | 0.1 | 85 | 94 | 73 | 100 | 100 | 82 | 100 | 100 | 100 |
| KIT (V559D) | | 46 | 4 | 0.15 | 97 | 100 | 91 | 100 | 100 | 78 | 100 | 100 | 65 |

The compounds were screened at concentrations of 0.1, 1.0 and10 mM , and results for primary screen binding interactions are reported as '% Ctrl', where lower numbers indicate stronger hits in the matrix. Red highlights those hits as <30% of Ctrl.

**Supplementary Table 6: KINOMEscan™ Selectivity Score for HC4, HC19, HC28 and GSK157 at concentrations of 0.1, 1.0 and 10 μM across 468 Kinases**

| Compound | Selectivity Score Type | Number of Hits | Number of Non-Mutant Kinases | Screening Concentrations | Selectivity Score |
|---|---|---|---|---|---|
| HC4 | S(35) | 0 | 403 | 100 | 0 |
| HC4 | S(10) | 0 | 403 | 100 | 0 |
| HC4 | S(1) | 0 | 403 | 100 | 0 |
| HC4 | S(35) | 2 | 403 | 1000 | 0.005 |
| HC4 | S(10) | 0 | 403 | 1000 | 0 |
| HC4 | S(1) | 0 | 403 | 1000 | 0 |
| HC4 | S(35) | 9 | 403 | 10000 | 0.022 |
| HC4 | S(10) | 3 | 403 | 10000 | 0.007 |
| HC4 | S(1) | 0 | 403 | 10000 | 0 |
| HC19 | S(35) | 0 | 403 | 100 | 0 |
| HC19 | S(10) | 0 | 403 | 100 | 0 |
| HC19 | S(1) | 0 | 403 | 100 | 0 |
| HC19 | S(35) | 0 | 403 | 1000 | 0 |
| HC19 | S(10) | 0 | 403 | 1000 | 0 |
| HC19 | S(1) | 0 | 403 | 1000 | 0 |
| HC19 | S(35) | 11 | 403 | 10000 | 0.027 |
| HC19 | S(10) | 4 | 403 | 10000 | 0.01 |
| HC19 | S(1) | 0 | 403 | 10000 | 0 |
| HC28 | S(35) | 1 | 403 | 100 | 0.002 |
| HC28 | S(10) | 0 | 403 | 100 | 0 |
| HC28 | S(1) | 0 | 403 | 100 | 0 |
| HC28 | S(35) | 3 | 403 | 1000 | 0.007 |
| HC28 | S(10) | 0 | 403 | 1000 | 0 |
| HC28 | S(1) | 0 | 403 | 1000 | 0 |
| HC28 | S(35) | 17 | 403 | 10000 | 0.042 |
| HC28 | S(10) | 6 | 403 | 10000 | 0.015 |
| HC28 | S(1) | 0 | 403 | 10000 | 0 |
| GSK157 | S(35) | 2 | 403 | 100 | 0.005 |
| GSK157 | S(10) | 0 | 403 | 100 | 0 |
| GSK157 | S(1) | 0 | 403 | 100 | 0 |
| GSK157 | S(35) | 21 | 403 | 1000 | 0.052 |
| GSK157 | S(10) | 7 | 403 | 1000 | 0.017 |
| GSK157 | S(1) | 1 | 403 | 1000 | 0.002 |
| GSK157 | S(35) | 93 | 403 | 10000 | 0.231 |
| GSK157 | S(10) | 39 | 403 | 10000 | 0.097 |
| GSK157 | S(1) | 9 | 403 | 10000 | 0.022 |

**Supplementary Table 7: Aqueous kinetic solubility, Caco-2 permeability, plasma protein binding and hepatocyte stability of HC4, HC19, HC28 and GSK-157**

| HC # | Aq Sol (µM) | Caco-2 $P_{app}$ ($X10^{-6}$ cm/s) | | Plasma Protein Binding, % | | | | Hepatocyte Cl ($t_{1/2}$ min) | | | |
|---|---|---|---|---|---|---|---|---|---|---|---|
| | | A-B | B-A | Human | Rat | Mouse | Dog | Human | Rat | Mouse | Dog |
| 5404 | 31 | 15 | 30.0 | 99.3 | 97.6 | 98.3 | 98.6 | 245 | 9.3 | 69 | 187 |
| 5490 | 48 | 14 | 45.7 | 98.1 | 96.7 | 97.2 | 97.1 | 118 | 34 | 69 | 877 |
| 5580 | 10 | 22 | | 99.7 | 99.3 | 99.4 | 99.8 | 310 | 14 | 63 | 493 |
| GSK157 | 27 | 6.1 | | 99.5 | 99.6 | 97.8 | 94.6 | | | | |

### Supplemental figure legends

**Supplementary Figure 1** (a) Flow diagram of the steps followed for single cell gene expression analysis with C1 and Biomark HD Fluidigm. A total of 255 DTCs and 90 primary tumor (PT) cells were analyzed. (b) List of the genes analyzed by high-throughput qPCR. (c) (upper) Immunoblot showing the inhibition of PERK phosphorylation (T982) by the PERK inhibitor HC4 (2 μM) in MCF10A-HER2 cells plated on low attachment plates and incubated with inhibitor for 24h. (lower) Immunoblot showing inhibition of PERK phosphorylation by treatment with increasing doses of HC4, HC19, HC28 and GSK2656157 for 4h upon incubation with ER stress inducer tunicamycin. (d) MCF10A cells were treated with DMSO, thapsigargin (0.2 μM) or PERK inhibitors (2 μM) in combinations as indicated in the figure for 24 h and assayed for GADD34 expression by qRT-PCR. N=3 wells per group. ****, p <0.0001. (e) The PERK inhibitor HC4 sensitizes to low dose ER stress-induced cell death in vitro. HC4 dose-response viability curve (Cell Titer Blue, CTB) in MCF10A-HER2 cells, in the absence (-) or in the presence of stress (low dose thapsigargin, Tg 2 nM) after 48 h (N=4) . Dashed line indicates IC50 (≈ 9 nM). (f) In vivo pharmacokinetic profile of HC4 in mouse plasma. Dashed lines indicate the different biochemical, cellular and in vitro viability $IC_{50}$s. (g) Effect of HC4 on total bone marrow cells (in two lower limbs) in MMTV-HER2 females treated for 2 weeks. Effect of HC4 on total white blood cells in MMTV-HER2 females treated for 2 weeks.

**Supplementary Figure 2.** (a) Normalized area of single macro-metastases in vehicle and HC4-treated animals (N=21 and 15). P by Mann-Whitney test. (b) Quantification of circulating tumor cells/ml blood by HER2 staining of cytospins (N=4). (c) Percentage P-Rb+ micro-mets per lung section/animal (N=4 and 6). P by Student's t test.

**Supplementary Figure 3.** (a) PERK (EIF2AK3) is upregulated in a sub-population of HER2+ breast cancer patients. Analysis of TCGA breast cancer data HER2+ cases (120 tumors) using cBioPortal. (b) Representative images of carmine staining of whole mount FVB normal mammary gland compared to vehicle- and HC4-treated MMTV-neu mammary gland whole mount. (c) Examples of the mammary gland histological structures quantified. Scale bars 100 and 25 μm.

**Supplementary Figure 4.** (a) Western blot for P-PERK levels in MMTV-neu tumor lysates from vehicle- and HC4-treated animals. (b) Tumor volumes from vehicle- (upper) and HC4-treated (lower) females ($mm^3$). Each line represents a tumor. (c) Percentage decrease in tumor size in HC4-treated females that showed tumor shrinkage. Each line represents a tumor and animal. (d) IHC for P-histone H3 in mammary gland tumor sections, representative images and quantification (right graph, N=5). P by Mann-Whitney test. (e) HER2-overexpressing ZR-75-1 cells were seeded on matrigel and after acinus establishment (day 4) wells were treated with vehicle (control) or HC4 (2 μM) for 10 days. Percentage of cleaved caspase-3 positive cells per acini (N=20) ± s.d. P by Student's t test. (f) MCF10A-HER2 cells were seeded on Matrigel and after acinus establishment (day 4) wells were treated with vehicle (control) or HC4 (2 μM) for 10 days. Percentage of P-histone H3-positive cells per acini (N=20) ± s.d. P by Student's t test.

**Supplementary Figure 5.** (a) Scoring system used for the quantification of P-HER2 levels in mammary gland tumor sections. The IHC P-HER2 positive area was multiplied by its intensity score according to established score shown in these representative images. Scale bar, 100 μm. (b) MCF10A-HER2 cells were starved o/n and treated +/-HC4 (2 μM), after which +/-EGF (100 ng/ml) was added for 20 min before collection. The levels of P-HER2/Y1112 and P-

HER2/Y877 were assessed by Western blot. GAPDH and HSP90 were used as loading controls. (c) Input for the extracts used in the surface biotinylation assay.

**Supplementary Methods**

**Kinetic Solubility Assay**. Test compound in 1% DMSO was incubated in phosphate buffer saline pH7.4 for 1h, followed by determination via UV/Vis absorbance of dissolved concentration against a calibration curve.

**Caco-2 Permeability Assay**. Caco-2 cells (clone cells (clone C2BBe1) were obtained from American Type Culture Collection (Manassas,VA). Cell monolayers were grown to confluence on collagen-coated, microporous membranes in 12-well assay plates. The permeability assay buffer was Hanks' balanced salt solution (HBSS) containing 10 mM HEPES and 15 mM glucose at a pH of 7.4. The buffer in the receiver chamber also contained 1% bovine serum albumin. The dosing solution concentration was 5 µM of test article in the assay buffer +/- 1 µM valspodar. Cells were first pre-incubated for 30 minutes with HBSS +/- 1 µM valspodar. Cell monolayers were dosed on the apical side (A-to-B) or basolateral side (B-to-A) and incubated at 37°C with 5% CO2 in a humidified incubator. Samples were taken from the donor and receiver chambers at 120 minutes. Each determination was performed in duplicate. The flux of lucifer yellow was also measured post-experimentally for each monolayer to ensure no damage was inflicted to the cell monolayers during the flux period. All samples were assayed by LC-MS/MS using electrospray ionization. The apparent permeability (Papp) was calculated as follows: $Papp=(dC_r / dt) \times V_r / (A \times C_A)$, where $dC_r / dt$ is the slope of the cumulative concentration in the receiver compartment vs time in $\mu M\ s^{-1}$; $V_r$ is the volume of the receiver compartment in $cm^3$; A is the area of the insert (1.13 $cm^2$ for 12-well); $C_A$ is the average of the nominal dosing concentration and the measured 120-minute donor concentration in µM.

**Protein Plasma Binding Assay**. Test compound was prepared at 1 μM concentration in incubation buffer containing 0.5% acetonitrile and incubated with plasma for 4h prior to LC-MS/MS compound determination. Protein binding of the test compound was measured using Rapid Equilibrium Dialysis (RED).

**Hepatocyte Intrinsic Clearance Assay**. Human hepatocytes pooled from 10 donors were incubated ($10^6$ viable cells/ml) with test compound at 1 μM concentration (0.1 % DMSO) and metabolic clearance determined by measuring percent remaining of test compound at multiple time points during incubation (15, 30, 60, 90 and 120 minutes). Compound concentration was determined by LC-MS/MS.

**KINOMEscan™ Kinase Selectivity Assay**. For most assays, kinase-tagged T7 phage strains were grown in parallel in 24-well blocks in an E. coli host derived from the BL21 strain. E. coli were grown to log-phase and infected with T7 phage from a frozen stock (multiplicity of infection = 0.4) and incubated with shaking at 32°C until lysis (90-150 minutes). The lysates were centrifuged (6,000 x g) and filtered (0.2μm) to remove cell debris. The remaining kinases were produced in HEK-293 cells and subsequently tagged with DNA for qPCR detection. Streptavidin-coated magnetic beads were treated with biotinylated small molecule ligands for 30 minutes at room temperature to generate affinity resins for kinase assays. The liganded beads were blocked with excess biotin and washed with blocking buffer (SeaBlock (Pierce), 1 % BSA, 0.05 % Tween 20, 1 mM DTT) to remove unbound ligand and to reduce non-specific phage binding. Binding reactions were assembled by combining kinases, liganded affinity beads, and test compounds in 1x binding buffer (20 % SeaBlock, 0.17x PBS, 0.05 % Tween 20, 6 mM DTT). Test compounds were prepared as 40x stocks in 100% DMSO and directly diluted into the assay. All reactions were performed in polypropylene 384-well plates in a final volume of 0.02 ml. The assay plates were incubated at room temperature with shaking for 1 hour and the affinity beads were washed with wash buffer (1x PBS, 0.05 % Tween 20). The beads were then re-suspended in elution buffer (1x PBS, 0.05 % Tween 20, 0.5 μM non-biotinylated affinity ligand) and incubated at room temperature with shaking for 30 minutes. The kinase concentration in the eluates was measured by qPCR. Compounds that bind the kinase active site and directly (sterically) or indirectly (allosterically) prevent kinase binding to the immobilized ligand, will reduce the amount of kinase captured on the solid support. Screening hits are identified by measuring the amount of kinase captured in test vs control samples and reported as “% Ctrl”, where lower numbers indicate stronger hits.

Selectivity Score or S-score is a quantitative measure of compound selectivity. It is calculated by dividing the number of kinases that compounds bind to by the total number of distinct kinases tested, excluding mutant variants.

S = Number of hits / Number of assays

This value can be calculated using %Ctrl as a potency threshold (below) and provides a quantitative method of describing compound selectivity to facilitate comparison of different compounds.

S(35) = (number of non-mutant kinases with %Ctrl <35)/(number of non-mutant kinases tested)

S(10) = (number of non-mutant kinases with %Ctrl <10)/(number of non-mutant kinases tested)

S(1) = (number of non-mutant kinases with %Ctrl <1)/(number of non-mutant kinases tested)

**In vivo Pharmacokinetic Study**. Female CD1 mice (6-8 weeks of age) (N=5) were dosed with vehicle (10% ethanol, 90% corn oil) or HC4 at 50 mg/kg once and blood samples collected into tubes containing K2-EDTA at different timepoints (15 min, 30 min, 1h, 2h, 4h, 8h, 12h and 24h). Plasma samples were prepared by centrifugation. The plasma concentration of test compound was determined by protein precipitation and LC-MS/MS.
